# MMP12 Knockout Prevent Weight and Muscle Loss Induced by Cancer Cachexia

**DOI:** 10.1101/2021.01.29.428838

**Authors:** Lingbi Jiang, Mingming Yang, Shihui He, Zhengyang Li, Haobin Li, Ting Niu, Dehuan Xie, Yan Mei, Xiaodong He, Lili Wei, Pinzhu Huang, Mingzhe Huang, Rongxin Zhang, Lijing Wang, Jiangchao Li

## Abstract

Weight loss and muscle wasting can have devastating impacts on survival and quality of life of patients with cancer cachexia. Here, we have established a hybrid mouse of Apc^Min/+^ mice and MMP12 knockout mice (Apc^Min/+^; MMP12^-/-^) and found that knockout MMP12 can suppress the weight and muscle loss of Apc^Min/+^ mice. In detail, we found that interleukin 6 was highly upregulated in the serum of cancer patients and MMP12 was increased in muscle of tumor-bearing mice. Interestingly, the interleukin 6 secreted by tumor cells led to MMP12 overexpression in the macrophages, which further resulted in degradation of insulin and insulin-like growth factor 1 and interruption of glycolipid metabolism. Notably, depletion of MMP12 prevented weight loss of Apc^Min/+^ mice. Our study uncovers the critical role of MMP12 in controlling weight and highlights the great potential of MMP12 in the treatment of cancer cachexia.

## 1. Introduction

Many studies have shown that rapid skeletal muscle mass loss is a characteristic of cancer cachexia (CAC) in colorectal cancer (CRC) patients with advanced cancer stages, which cannot be completely reversed by conventional nutritional support or drugs therapy (Bonetto et al., 2016; Pettersen et al., 2017; Song et al., 2019; Yang et al., 2018; Yang et al., 2019; Yuan et al., 2015). Muscle loss caused by tumor development or growth would occur in various cancers, such as pancreatic cancer, esophageal cancer, gastric cancer, lung cancer, liver cancer and CRC. According to the statistics, nearly 80% of cancer patients have skeletal muscle loss as a late outcome, and the mortality rate is as high as 30%(Baracos et al., 2018; Herremans et al., 2019; Tisdale and Michael, 2002). 2018 estimates of global cancer data show that CRC is the third most common malignant tumor in the world, and its incidence and mortality are increasing year by year (Bray et al., 2018; Brody, 2015; Lin et al., 2016; Siegel et al., 2014), and cachexia muscle loss induced by CRC is an important cause of death. Although more and more attention has been paid to CAC in recent years and great progress has been made in the diagnosis and treatment of CRC, the mortality rate of CRC has not decreased and has been younger (Brody, 2015; Center et al., 2009). The key is that the mechanism of how the inflammatory environment of tumor causes muscle loss is still not clear, and CAC involves a variety of immune cells, a variety of cytokines, and metabolic disorders(Daou, 2020). The tumor-induced pro-inflammatory response plays an important role in the progression of CAC. This metabolic dysfunction is caused by changes in glucose, lipid and protein metabolism, and may lead to the loss of skeletal muscle and adipose tissue when maintained for a long time (Fonseca et al., 2020; Lobato et al., 2018; Patel and Patel, 2017). To date, although several drugs have had positive clinical effects in increasing lean body mass, their effects on body function are limited. There are no effective medical interventions or approved drug therapies that can completely reverse muscle loss caused by CAC, which brings difficulties to the treatment of chemotherapy drugs (Daou, 2020) (Fonseca et al., 2020).

MMP12 is a matrix protein metalloenzyme, also named macrophage metalloenzyme, from a family of endoproteolytic enzymes whose activities depend on metal ions such as calcium and zinc and can degrade extracellular matrix. It was discovered in a study of tadpole morphological changes during development and is necessary for monocyte recruitment, and it is mainly secreted by inflammatory cells such as monocyte macrophages(Bauters et al., 2013). MMP12 is mainly secreted by M2 macrophages(Han et al., 2018; Hotary et al.; Lee et al., 2016). Reports have demonstrated that MMP12 can decompose most extracellular matrix and vascular components, and has obvious effects on elastic fiber-rich blood vessels, lung, embryonic development, reproduction, and tissue remodeling(Atlı, 2017; Kraen et al., 2019; Langlois et al.; Wagner et al., 2016; Wang et al., 2019; Wetzl et al.). MMP12 is associated not only with smoking-induced emphysema(Kraen et al., 2019) but also with the typing of bone marrow cells and myeloid derived suppressor cells(Qu et al., 2011). As early as 1981, studies reported that MMP12 can specifically degrade insulin(Kettner et al.). In 2014, researchers of Washington University confirmed that MMP12 regulates insulin sensitivity and is positively correlated with insulin resistance(Lee et al., 2016). In 2016, MMP12 was identified as a target for insulin-related treatment of metabolic diseases, and MMP12 promotes insulin resistance and prevents fat expansion under high-fat conditions(Amor and Moreno-Viedma; Bauters et al., 2013).

In recent years, some inflammatory cytokines, such as Interleukin 6 (IL-6), monocyte chemoattractant protein-1, tumor necrosis factor, zinc-α2-glycoprotein and pancreatic enzymes, have been shown to be related to muscle loss (Bing, 2011; Han et al., 2018; Pettersen et al., 2017; Talbert et al., 2018a; Yarla et al., 2018). IL-6, a multi effect proinflammatory cytokine, is secreted by normal human monocytes, fibroblasts, endothelial cells, Th2 cells, vascular endothelial cells. And a variety of tumor cells also secrete IL-6 (Carson and Baltgalvis, 2010b; Han et al., 2018; Pettersen et al., 2017). It targets macrophages, hepatocytes, resting T cells, activated B cells and plasma cells (Han et al., 2018; Utsumi et al., 1990). The IL-6 level has been proposed to be high in patients with skeletal muscle loss (Peixoto da Silva et al., 2020a). IL-6 levels were increased in tumor tissue and involved in skeletal loss progression in cancer patient (Narsale et al., 2014). IL-6 can directly induce alternative macrophage activation(Ayaub et al.; Hopkins et al.). And systemic overexpression of IL-6 accelerates CAC muscle loss in Apc^Min/+^ mice (Baltgalvis et al., 2010). A high level of IL-6 in serum is considered to be an important contributor to the progression of muscular dystrophies, including muscle loss induced by CAC and duchenne muscular dystrophy. Blocking IL-6 receptor may inhibit dystrophic muscle loss and lipolysis by suppressing the downstream Janus kinase/signal transducer and transcription activator (JAK/STAT) pathway to promote muscle regeneration(Hu et al., 2019; Wada et al., 2017). Ville Wallenius et al found that centrally acting IL-6 exerts anti-obesity effects in rodents but does not increase energy expenditure(Franckhauser et al., 2008; Wallenius et al., 2002). IL-6 has been shown to induce insulin resistance(Liaqat et al., 2017). It is well known that insulin affects glucose uptake through the PI3K-AKT-mTOR pathway by binding to insulin receptors(Hopkins et al.). Skeletal muscle is the primary tissue involved in insulin-stimulated glucose uptake. IL-6 mediates glucose intolerance and promotes insulin resistance in skeletal muscle (Deshmukh et al.; Han et al., 2018; Nicholson et al., 2019). Moreover, insulin resistance can promote muscle wasting(Peixoto da Silva et al., 2020b). IL-6 suppresses insulin action through the Signal transducer and activator of transcription 3 (STAT3) pathway, which then may affect the insulin receptor by suppressing insulin receptor substrate-1 and downstream targets(Puppa et al., 2012). In short, various studies have proven that IL-6 can induce insulin resistance, thereby indirectly exacerbating muscle loss in CAC. In our study, we found that IL-6 secreted by tumors would target muscle macrophages, and then MMP12 in these activated macrophages will be upregulated, degrading insulin and insulin-like growth factor 1. So, we thought one molecular crosstalk may exist in bearing mice. Increased expression of IL-6 derived from cancer cells up-regulates MMP12 in macrophages, which affects skeletal glycolipid metabolism over a long period of time, may resulting in loss of skeletal muscle for a long time.

To date, for patients with muscle and weight loss induced by CAC, multimodal interventions including drugs, nutritional support and physical exercise may be a reasonable approach for future research to better understand and prevent loss of muscle (Fonseca et al., 2020). Although several drugs have had positive clinical effects in increasing lean body mass, their effects on body function are limited. There are no effective medical interventions or approved drug therapies that can completely reverse muscle loss caused by CAC, which brings difficulties to the treatment of chemotherapy drugs (Daou, 2020) (Fonseca et al., 2020). Taken together, the combination of MMP12 inhibitors and chemotherapy drugs may bring new challenges and ideas for the treatment of cancer cachexia to improve the quality of life.

## 2. Materials and Methods

### 2. 1 Mice

B6.129X-MMP12tm1Sds/J macrophage metalloelastase-deficient (MMP12^-/-^) mice (no. 004855), with a C57BL/6 background were purchased from the Jackson Laboratory, USA(https://www.jax.org/strain/004855). Apc^Min/+^ mice (no. T001457) were obtained from Gem Pharmatech, China (http://www.gempharmatech.com/cn/index.php/searchinfo/59/17.html). Wild-type (WT/ C57BL/6J) mice were purchased from Guangdong Medical Laboratory Animal Center (China). The production license number was SCXK (Guangdong) 2017-0125. Apc^Min/+^; MMP12^-/-^ mice were obtained by crossing Apc^Min/+^ mice and MMP12^-/-^ mice (Figure 1-figure Supplement 1A). All mice were housed in specific pathogen–free conditions. All mice studies complied with the Guangdong Pharmaceutical University, and all protocol was approved by the animal experimental ethics committee of Guangdong Pharmaceutical University.

### 2. 2 Genotype identification

We established crossbred mice and genotyped the 3-week-old mice. The polymerase chain reaction products were subjected to gel electrophoresis (1.2%), and a gel imaging system (GboxGyngene system, UK) was used to obtain electrophoresis images (Figure 1-figure Supplement 1B). The details of genotype identification can be found on the website of the Jackson Laboratory https://www.jax.org/strain/004855.

### 2. 3 Mice experiments and Tissue collection

Mice were anesthetized by inhalation of carbon dioxide anesthesia, marrow, blood, epididymal white fat, gastrocnemius and soleus muscle and brown back fat were collected. For immunohistochemistry Mice were anesthetized by inhalation of carbon dioxide anesthesia, marrow, blood, epididymal white fat, gastrocnemius and soleus muscle and brown back fat were collected. For immunohistochemistry staining analysis, we obtained muscle tissues wax tissue blocks from clinical surgery patients. For serum detection, all fresh clinical blood samples were obtained from the Sun Yat-Sen University Cancer Center, Guangzhou, China. all fresh clinical blood samples were taken individuals undergoing a clinical health individuals and colorectal cancer patients (30-60 years, excluding individuals with diabetes and hyper-thyroidism) and frozen at −80℃ until the experiments were performed.

### 2. 4 Antibodies and Reagents

An anti-F4/80 anti-body (Cat: 14-4801-81) was purchased from eBioscience and anti-MMP12 (MA5-24851) were purchased from Thermo Fisher (Thermo Fisher Scientific, Cambridge, Massachusetts, USA). An anti-GAPDH (5174P) and anti-β-actin (4970S) were purchased from Cell Signaling Technology Inc (CST). Recombinant mouse MMP-12 protein (3467-MPB-020) was purchased from R&D Systems, Inc. The MMP12 inhibitor MMP408 (444291) was purchased from Merck Millipore Company. Alexa Fluor-488 donkey antibody (P/N SA11055S) was purchased from Invitrogen (Thermo Fisher Scientific, Cambridge, Massachusetts, USA).

### 2. 5 Total RNA extraction and Real-time polymerase chain reaction

All tissues from mice were quick-frozen in liquid nitrogen and stored at −80°C until they were dissolved with Trizol (TaKaRa, Guangzhou, China, A161050A). RNA extraction was performed according to the manufacturer’s instruction, and the total extracted RNA was reverse-transcribed into cDNA for polymerase chain reaction amplification using the real-time polymerase chain reaction SYBR Green kit (TaKaRa, Guangzhou, China). The reverse transcription steps were as follows: denaturation at 94°C for 5 minutes; 40 cycles of denaturation at 94°C for 30 seconds, annealing at 60°C for 30 seconds, and extension at 72°C for 30 seconds; 72°C for 5 minutes. The mRNA samples were quantified in triplicate. The house-keeping gene GAPDH was used as an internal control to normalize the real-time polymerase chain reaction data for each sample of mRNA. All real-time polymerase chain reaction primers were synthesized by Shanghai Sangon Biotechnology Inc., China, and the primer sequences are listed in S Table 1.

**Table1:**
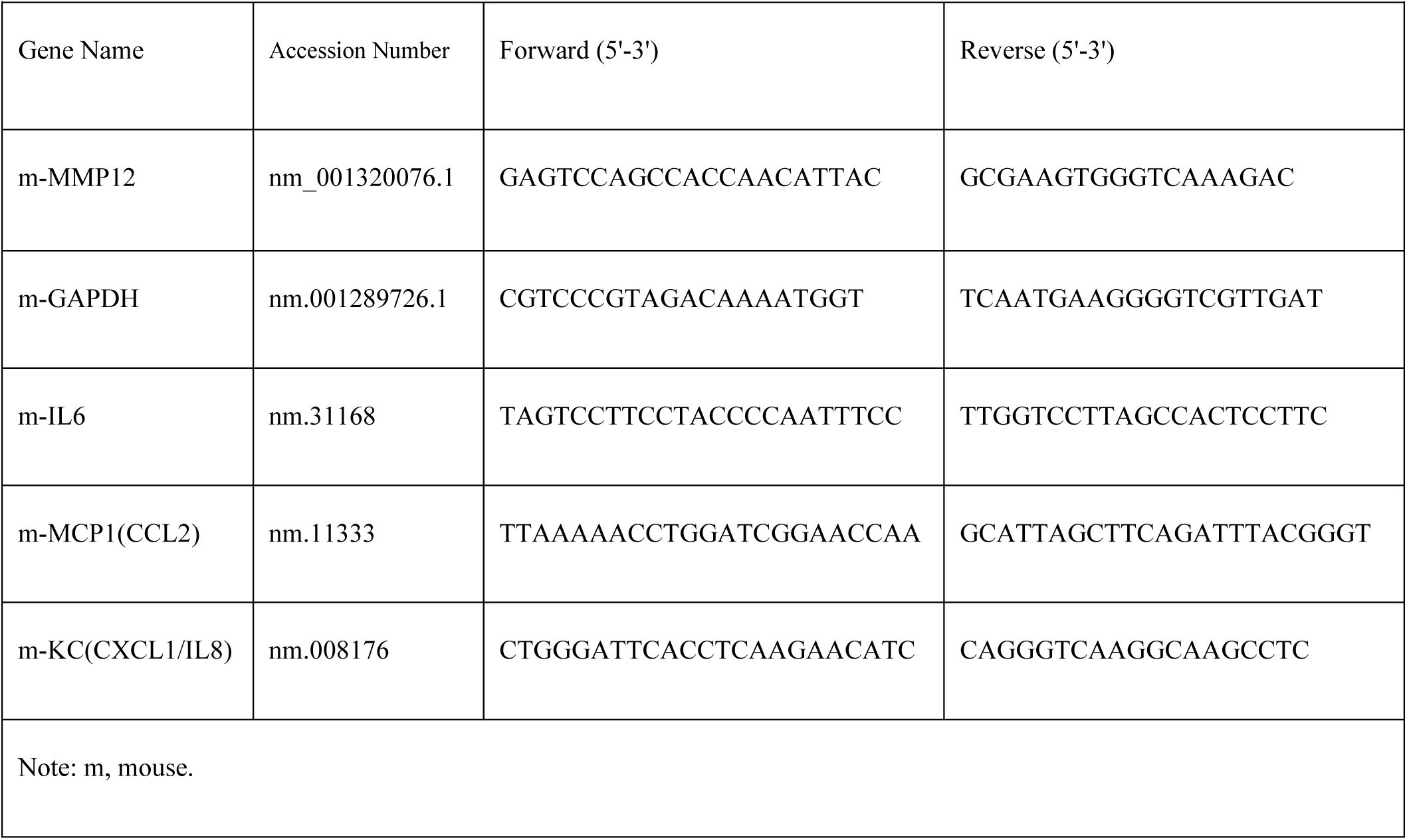
Quantitative PCR primers.

### 2. 6 Histological analysis and Hematoxylin and eosin staining

Formalin-fixed (10% neutral buffered formalin), gradient dehydration and paraffin-embedded to obtain 3-μm tissue sections from mice were subjected to perform the experiments of hematoxylin and eosin staining, immunohistochemical staining and immunofluorescence staining in accordance with the protocols. To assess the cross-sectional area of muscle, image J software was used after hematoxylin and eosin staining

### 2. 7 Immunohistochemistry

Tissue sections were dewaxed and incubated with 30% hydrogen peroxide in methanol and blocked with 10% bovine serum albumin diluted with phosphate buffered saline (PBS). The sections were incubated with primary antibodies against MMP12 (1:100) at 4°C overnight. Finally, the primary antibody-treated sections were incubated with secondary antibody (1:100) horseradish peroxides (HRP) (goat anti-rabbit IgG) conjugated with HRP at 37°C for 1hour, stained with 3,3-diaminobenzidine, and counterstained with hematoxylin.

### 2. 8 Double immunofluorescence staining

Tissue sections were dewaxed and blocked with 10% bovine serum albumin diluted with PBS. The sections were incubated with a mixture of primary antibodies against MMP12(1:100) and F4/80 (1:100) overnight at 4°C. The next day, the primary antibody-treated sections were incubated with mixtures of secondary antibodies conjugated Alexa Fluor 488 (1:100) and Alexa Fluor 555 (1:100) for 1 hour at room temperature. Immunostaining signals and DAPI-stained nuclei were visualized under a confocal microscope.

### 2. 9 Enzyme-linked immunosorbent assay

The enzyme-linked immunosorbent assay (ELISA) was performed with kits with serum samples from patients and mice according to the manufacturer’s protocol. The human-IL6 kit (EHC007), mouse JE/MCP1/CCL2 (EMC113) kit, mouse IL-6 ELISA kit (EMC004), and mouse KC/IL-8/CXCL1 ELISA kit (EMC104) were purchased from NeoBioscience Technology Company (ShenZhen, China). The rat/mouse insulin kit (EZRMI-13K) was purchased from EMD Millipore Corporation. A mouse MMP12 ELISA kit (ARG81803) was purchased from Arigobiolaboratories company. The human CXCL1/KC kit (EK-196) was purchased from Multi Science Company. The data of ELISA was analyzed by using Curve Expert1.4 software.

### 2. 10 Western blotting

Tissue samples (50-80 mg) or cells were homogenized and lysed with radio immunoprecipitation assay buffer (Thermo Scientific, 89900) containing with protease and phosphatase inhibitors, and then the supernatants were clarified by centrifugation. Quantitative analysis based on the bicinchoninic acid (BCA) protein assay was used to detect the protein concentration. Denatured proteins in the supernatant were separated by sodium dodecyl sulfate–polyacrylamide gel electrophoresis and transferred to polyvinylidene difluoride (Millipore Corporation, Billerica, MA, USA) membranes, blocked with 5% nonfat milk powder at room temperature, and then incubated with the primary antibodies (1:1000) overnight at 4°C. The next day, the protein strips were further incubated with HRP-conjugated anti-rabbit secondary antibodies (1:5000) and the bands were visualized after exposure to film after incubated with enhanced chemiluminescence detection reagents. These bands were visualized after exposure to film. We used imageJ software to analyze the optical density of the protein band. All experiments were repeated three times.

### 2. 11 Cytokine antibody assay

Qualitative assessment of 38 cytokines in the supernatants from media for culture (+MC38) or non-culture with MC38 (-MC38) cells was performed with the Ray Bio Mouse Cytokine Antibody Array 5 (AAM-INF-1-2, Ray Biotech) according to the provided manufacturer’s protocol. The detection procedure was as follows: The membranes were blocked by incubation with the blocking buffer. Diluted biotin-conjugated anti-cytokine antibodies and HRP-conjugated streptavidin were detected to immuno-complexes. The visualized X-ray film was exposed to chemiluminescence for quantification with ECL chemiluminescence. Semiquantitative data analysis was performed for signal intensity ImageJ, and positive controls were used to normalize the results. Every cytokine to positive control ratio (cytokine density/positive control density) is used to represent the relative content of every cytokine. The cytokines and their abbreviations are shown in (Figure 3-figure Supplement 7).

### 2. 12 Oral glucose tolerance test, Insulin tolerance test and Blood glucose level measurement

Mice were fasted for 8 hours and then housed overnight. And then, they were given either oral glucose (2 g/kg body weight or an intraperitoneal insulin injection (0. 75 IU/kg). The tail vein blood glucose level of mice was measured for tail blood glucose at 0, 15, 30, 45, 60, 90, and 120 minutes after treatment.

Blood samples were collected at 0, 15, 30 and 60, 90, and 120 minutes for glucose measurement in tail vein blood with a blood glucose meter (Johnson & Johnson) at the specified time points. All blood glucose levels were performed using the glucose meters.

### 2. 13 Serum lipid composition assay

The levels of total cholesterol (TC), total triglycerides (TG), high density lipoprotein cholesterol (HDL-C) and low density lipoprotein cholesterol (LDL-C) were determined according to the manufacturer’s protocols. The assay kits all were purchased from Jiancheng (Nanjing, China).

### 2. 14 Cell culture

The RAW264.7 cell line, MC38 cell line and CT26 cell line were purchased from the American Typical Culture Collection (ATCC) and cultured according to international standard protocols. All cell lines were maintained in Dulbecco’s Modified Eagle’s Medium (DMEM, Thermo Scientific HyClone, Beijing, China) +10% fetal bovine serum (FBS, HyClone) + 1% penicillin (HyClone)/streptomycin (HyClone). The RAW 264.7 cell line, MC 38 cell line and CT26 cell line were cultured in DMEM. All cell lines in the experiments were incubated with a mixture of 95% air and 5% CO_2_.

### 2. 15 Co-culture experiment

All cells were grown in DMEM+10% FBS+10% penicillin-streptomycin. The co-culture of RAW264.7 and MC38/CT26 cells was seeded into performed using a chamber with filter inserts (pore size,0.4 µm) in 6-well dishes (Corning, NY, USA). All cell lines could not pass to the filter because the pore size of the filter was smaller than the diameter of cell lines. RAW264.7 cells non co-cultured with MC38/CT26 cell lines (-MC38/CT26) were used as the negative controls. MC38/CT26 cell lines (control, 1×10^4,^ 3×10^4^, 5×10^4^) were seed in upper chamber, while RAW264.7cells (1-2×10^5^) were seed in the lower chamber. We can separate physically RAW264.7 cells or MC38/CT26 cell lines to obtain RAW264.7 (+MC38/CT26) cells in lower chamber. The RAW264.7 cells were homogenized and lysed with radio immunoprecipitation assay buffer to quantitative analysis and then subjected to western bloting.

### 2. 16 Interleukin 6 treatment of macrophages

The Interleukin 6 (IL-6) freeze-dried powder was purchased from Pepertech (216-16) and was dissolved in trehalose-bovine serum albumin aqueous solution. RAW264.7 cells (1-2×10^5^) were seeded into 6-well plates and treated with increasing different doses of IL-6 (+IL-6, 0, 2, 5 10,30ng/ ml) for 72hours. Next, RAW264.7 cells (1-2×10^5^) were treated continuously with IL-6(+IL-6, 30 ng/ml) for 0, 3, 6, and 9hours. Cells incubated with fresh media were used as the untreated (-IL-6) negative controls. Finally, western blotting was used to quantify MMP12 in RAW264.7 cells under different conditions.

### 2. 18 Isolation of primary peritoneal macrophages

24-week-old WT and Apc^Min/+^ mice were sterilized with 75% ethanol after cervical dislocation. The mouse abdomen was opened from the peritoneum, and 5 mL fatal bovine serum was injected with a syringe, which was allowed to remain inside the abdomen for 5 minutes, with gentle massaging for 30 s. The peritoneal fluid was collected, and this fluid was transferred to a 15 mL sterile tube to obtain peritoneal macrophages. After centrifugation (4°C, 1000 rpm) for 10 minutes, the supernatant was removed, and the collected cells were resuspended in Dulbecco’s Modified Eagle’s Medium. The resuspended cells were cultured in a petri dish for 2 hours (37°C), and the primary peritoneal macrophages were prepared from the remaining adherent cells after the medium was removed.

### 2.19 MMP12 and Peptide experiments

Recombinant mouse MMP-12 protein (3467-MPB-020) was purchased from R&D Systems, Inc. According to the instructions, dissolve MMP12 in a buffer containing 50mM Tris, 10 mM CaCl_2_, 150 mM NaCl, 0.05% (w/v) Brij-35, 5 µM ZnCl_2_, pH 7.5 at a concentration of 250ug/ml. Insulin polypeptide and insulin-like factor polypeptide were synthesized by ChinaPeptides Co., Ltd.

The fluorescent peptide sequence:

(1) Insulin:5-FAM-NQHLCGSHLVEALYLVCGERGFFYTPK(Dabcyl);

(2) insulin-like growth factor 1: 5-FAM-GPETLCGAELVDALQFVCGDRGFYFNK(Dabcyl). According to the instructions, the peptide freeze-dried powder was dissolved in 25% ACN and 75% ddH_2_O solvent at a concentration of 1 mg/ml.

The two experiments are as follows: (1) Fluorescence intensity: After mixed incubation of MMP12 and peptide (37℃, 2hours), the fluorescence intensity was measured with a fluorescence microplate reader.

(2) MS Analysis Report: After the MMP12 and peptide were mixed and incubated, the lower liquid after being filtered by a 34KD filter is subjected to Electrospray Ionization Mass Spectrometry (IMS) to detect its characteristic peaks. The experimental conditions are: Ion Source: ESI, Capillary (KV): ± (2500∼3000), Desolvation(L/hr):800, Desolvation Temp:450℃, Cone(V): 30∼50, Run Time: 1min.

### 2. 20 Mice administration

All 17-week-old Apc^Min/+^ mice were randomly divided into three groups (five mice in each group). A group of Apc^Min/+^ mice were administered intragastric with MMP12 inhibitor (MMP408) at a dose of 5mg/kg, and the other group was intraperitoneally administered with 5-FU (30mg/kg) combined with a dose of intragastric administration of MMP408. At the same time, Apc^Min/+^ mice injected with normal saline intraperitoneally served as a control group. The body weight was weighed by the administration every 2 days and after this continued for 10 days, the final weight change from the initial body weight was calculated.

### 2. 21 Data calculation and Statistical analysis

All mouse organ ratios represented the percentage of the organs/tissues weight compared to the body weight. The skeletal muscle to weight ratio was (Gastrocnemius +Soleus muscle)/body (%). All data were processed with GraphPad Prism 8.0 software and are presented as the means ± standard deviation (SD). A two-tailed test was used. Statistically significant differences were set at \**P*< 0.05, \*\**P*< 0.01, \*\*\**P*< 0.001, \*\*\*\**P*< 0.0001. All schematic images were created with BioRender.com.

## 3 Results

### 3. 1 Knockout MMP12 can suppress weight and muscle loss in Apc^Min/+^ mice

To investigate the weight dynamics of the mice, we determined the mouse body weight by retroactive examination of the body weight from 5-week to 24-weeks old. The current weight curves have shown that compared with the weight gain in wild type (WT) mice over time, the weight of Apc^Min/+^ mice reached its maximum peak at 12-week-old and then declined until to die at approximately 24-week-old (Figure 1A). Surprisingly, in comparison to the Apc^Min/+^ mice control, the body weight of Apc^Min/+^; MMP12^-/-^ mice increased by approximately 70% at the same age (Figure 1B). While, there were no significant differences between WT mice and MMP12^-/-^ mice in body weight (Figure 1C). As well known, the Apc^Min/+^ mouse is a model of muscle loss with cancer cachexia (CAC) and intestinal tumor burden. Because weight loss induced by CAC may be related to the wasting of skeletal muscle weight and fat weight (Peixoto da Silva et al., 2020a), here, we verified whether the weight gain of Apc^Min/+^mice caused by MMP12 knockout was due to reduction of fat and skeletal muscle loss at 24-week-old CAC stage in Apc^Min/+^ mice. We assessed the histologic white adipose tissue (WAT), compared with that in WT mice, the WAT-to-body weight ratio in Apc^Min/+^ mice decreased and the weight ratio of Apc^Min/+^; MMP12^-/-^ mice tended to increase compared with that of Apc^Min/+^ mice but the increase was not statistically significant (Figure 1D). However, a significant increase of approximately 4.5% in the muscle-to-body weight ratio was observed in Apc^Min/+^; MMP12^-/-^ mice compared with Apc^Min/+^ mice (Figure 1E).

**Figure 1.**
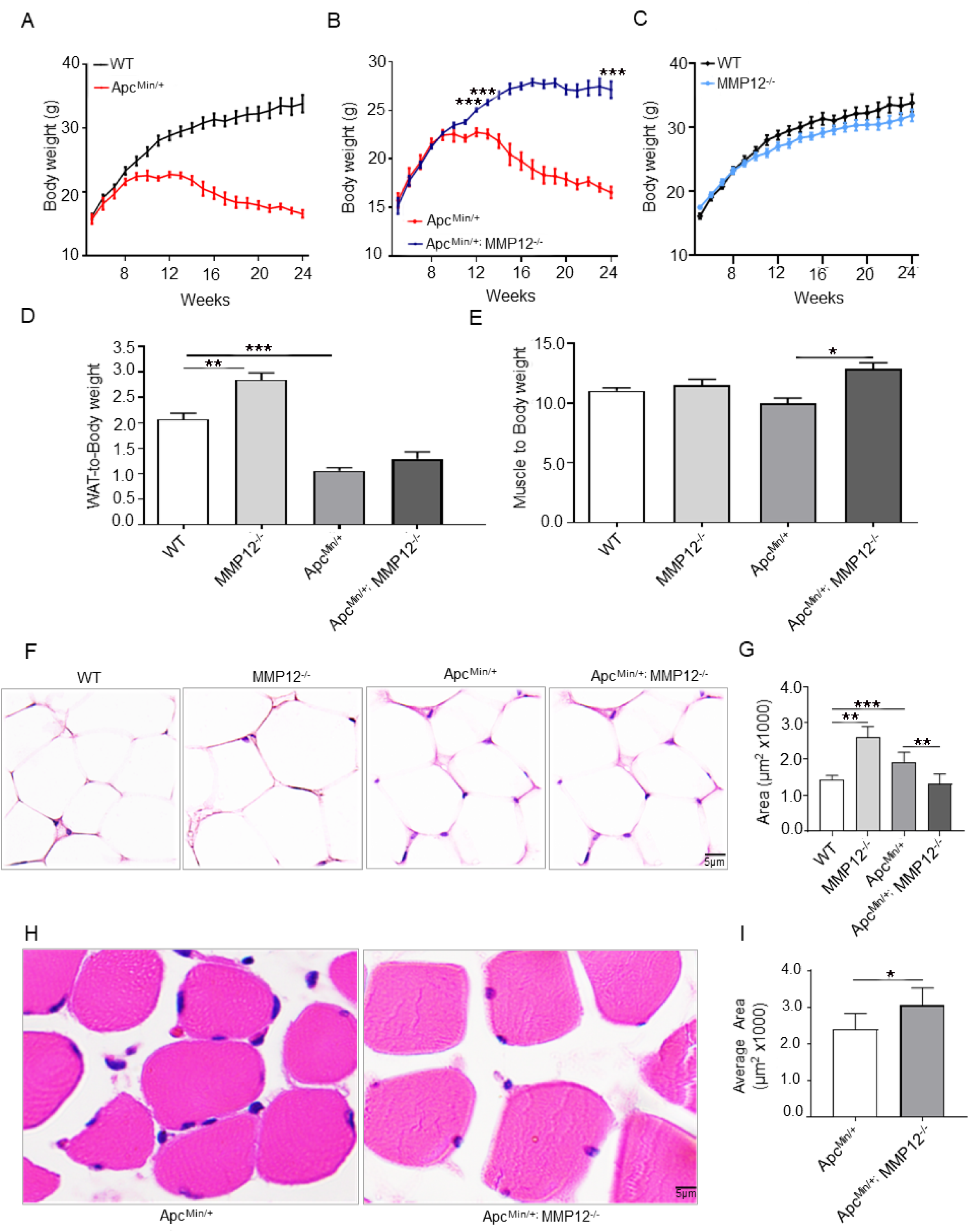
Knockout of MMP12 in Apc^Min/+^ mice prevents weight and muscle loss. (A-C) Plots of the body weight of wild-type (WT), Apc^Min/+^, Apc^Min/+^; MMP12^-/-^ and MMP12^-/-^ mice from 5 to 24 weeks (\*\*\**P*<0.001; \*\**P* < 0.01; \**P* < 0.05; data are shown as the means ± SD; n = 6 per group). (D) The ratio of white adipose tissue to body weight (\*\*\**P*<0.001; \*\**P* < 0.01, n=5). (E) The ratio of skeletal muscle to body weight (\**P<* 0.05, n=5). (F) Hematoxylin and eosin staining of white adipose tissue in WT, Apc^Min/+^, Apc^Min/+^; MMP12^-/-^ and MMP12^-/-^ mice at 24 weeks of age. Scale bars, 5μm. (G) Evaluation of the white adipose tissue across 4 groups by ImageJ software(40X) (\*\*\**P*<0.001; \*\**P* < 0.01, data are shown as the means ± SD; n = 5 mice each group). (H) Hematoxylin and eosin staining of muscle in Apc^Min/+^, Apc^Min/+^; MMP12^-/-^ at 24 weeks of age. Scale bars, 5μm. (I) Evaluation of the cross-sectional area of the gastrocnemius from Apc^Min/+^ mice and Apc^Min/+^;MMP12^-/-^mice by ImageJ software (40X) (\**P* < 0.05; data are shown as the means ± SD; n = 3 mice each group).

To further confirm the histological changes of WAT (Figure 1F) and muscle area (Figure 1H) in the four mice group at 24-week-old, we performed hematoxylin and eosin staining to assess the histological area by the ImageJ software (Figure 1F, H). We observed the fat area is larger in MMP12^-/-^ mice compared with WT mice, but there has no difference between Apc^Min/+^ mice and Apc^Min/+^; MMP12^-/-^ mice (Figure 1G). The H&E staining of muscle to show that the area of Apc^Min/+^; MMP12^-/-^ mice is estimated to be approximately 1-2-fold larger than Apc^Min/+^ mice (Figure 1I). Meanwhile, no difference in food intake was observed between Apc^Min/+^ and Apc^Min/+^; MMP12^-/-^ mice (Figure 1-figure Supplement 3A). Taken together, knocking out MMP12 prevents muscle from wasting in tumor-burden (Apc^Min/+^) mice at CAC stage, but not in WT mice.

### 3. 2 MMP12 is upregulated in muscle tissue and peritoneal macrophages of Apc^Min/+^ mice

To confirm whether MMP12 is expressed in muscle tissue, we use immunohistochemical staining, immunofluorescence staining and western immunoblotting to detect the expression of MMP12 in muscle. The immunohistochemistry results proved that MMP12 positive staining was expressed not only in skeletal muscles from clinical individuals (Figure 2A, Figure 2 -figure Supplement 3B), but also in WT mice (Figure 2B). In order to investigate why the reduction in muscle loss caused by knocking out MMP12 only occurred after tumor-bearing mice and not in WT mice, furthermore, we used immunohistochemistry methods to detect MMP12 expression in muscle from 24-week-old WT mice and Apc^Min/+^ mice (Figure 2C). The results of immunohistochemical staining showed that MMP12-positive staining was increased in the muscle of Apc^Min/+^ mice compared with that in WT mice at 24-weeks old by image J (Figure 2D). Because MMP12 is mainly secreted by macrophages (Lee et al., 2014a; Lee et al., 2014b), next, we performed double immunofluorescence (IF) to detect the expression of macrophages and MMP12 in muscle and found that the F4/80 and MMP12 markers were colocalized in mice (Figure 2E). Quantitative PCR (qPCR) revealed that a tendency towards higher MMP12 mRNA levels in peritoneal macrophages (as described in the Materials) was seen in Apc^Min/+^ mice (Figure 2F), which was consistent with immunohistochemistry results. We detected the dynamic circulating serum MMP12 level by enzyme-linked immunosorbent assay and found that the MMP12 levels in 9-, 15-, and 24-week-old WT mice and Apc^Min/+^ mice did not differ (Figure 2G). Taken together, MMP12 is expressed in muscle tissue and co-localized with macrophage. In comparison to the WT mice control, MMP12 is increased in skeletal muscle tissue and peritoneal macrophages of Apc^Min/+^ mice, but no difference in serum was witnessed between the two groups.

**Figure 2.**
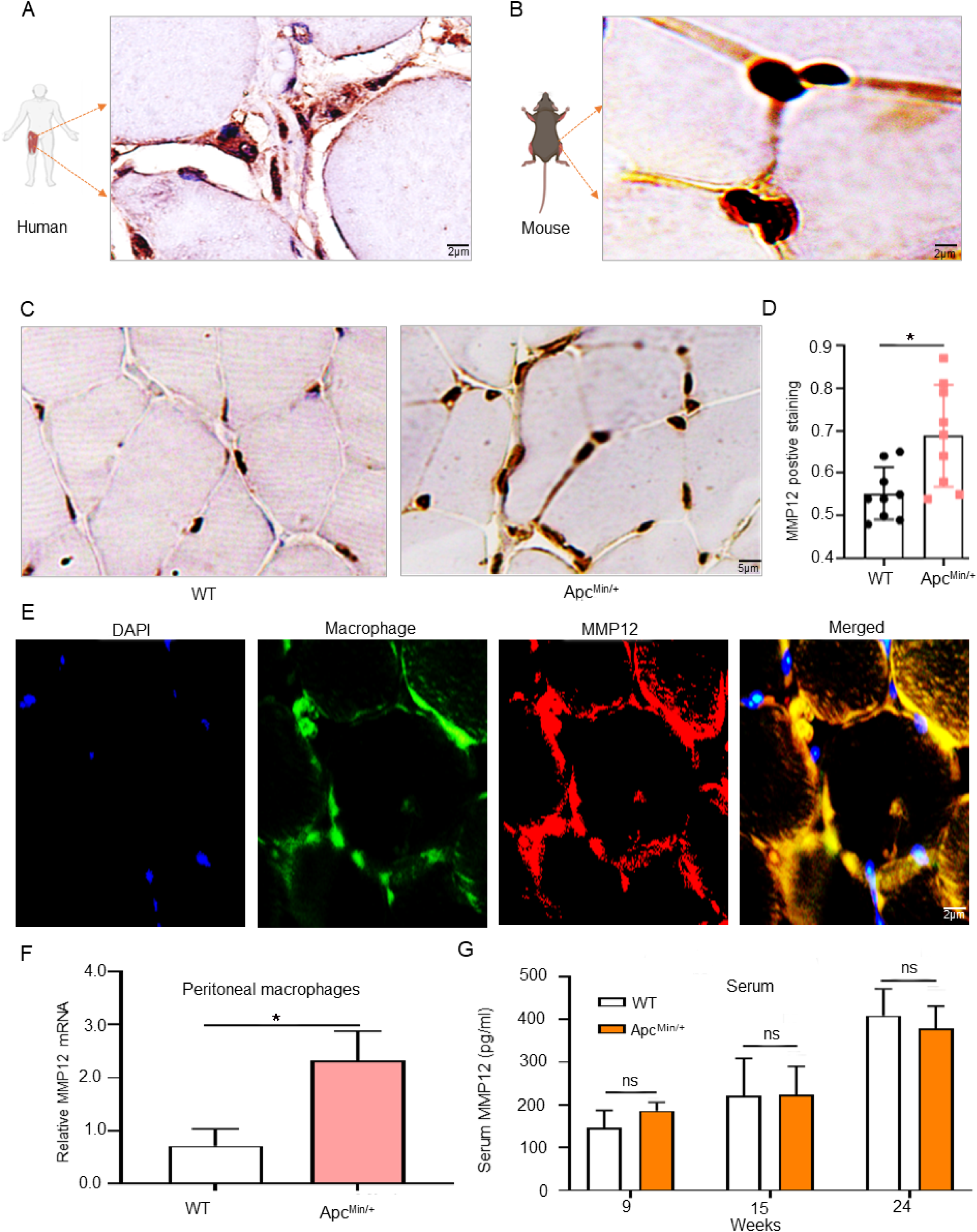
MMP12 was upregulated in muscle tissues and macrophages of Apc^Min/+^ mice. (A) Immunostaining of MMP12-positive in muscle of clinical individuals and WT mice. Scale bar, 2μm. (B) Immunostaining of MMP12-positive in muscle tissue of WT mice. Scale bar, 2μm (C) Immunostaining of MMP12-positive in muscle (gastrocnemius) from WT mice and Apc^Min/+^ mice at 24 weeks of age. Scale bar, 5μm. (D) Quantification of MM12-positive in gastrocnemius tissues was performed by ImageJ software (40X) (\**P<* 0.05, data are shown as the means ± SD; n = 4 per group). (E) Representative images of double immunofluorescent staining of macrophages (F4/80 in green) and MMP12 (in red) in WT mice are shown. The yellow areas in the merged images indicate overlapping localization of the red and green signals, indicated by the white arrows. Scale bars, 2μm. (F) Quantification of MMP12 mRNA expression level in peritoneal macrophages isolated from WT mice and Apc^Min/+^ mice by qPCR (\**P<* 0.05; data are shown as the means ± SD; n = 3 per group). (G) The serum MMP12 levels detected in WT and Apc^Min/+^ mice at 9-,15-, and 24 weeks by enzyme-linked immunosorbent assay (*P*> 0.05; data are shown as the means ± SD; n = 6 per group).

### 3. 3 Tumor cells can secrete IL-6

Previous studies have shown that interleukin 6 (IL-6) is one of the cytokines predictive muscle loss induced by CAC(Bonetto et al., 2012; Kim et al., 2013; Mahadik and Sujata, 2013; Mauer et al., 2014), it can accelerate muscle loss indued by CAC. And tumor cells are an important source of IL-6(Carson and Baltgalvis, 2010b; Han et al., 2018; Pettersen et al., 2017). The clinical literature data also suggested that among many CAC-muscle loss patients who lost weight and were close to death, IL-6 was almost the only increased cytokine among many factors. Therefore, we mainly focus on whether IL-6, which is related to muscle loss, is caused by tumors (Carson and Baltgalvis, 2010b). We observed that the clinical colorectal cancer patients had significantly higher serum IL-6 levels than the normal healthy group (Figure 3A). In vivo, a similar trend was found in Apc^Min/+^ mice, and serum IL-6 levels in Apc^Min/+^ mice were significantly increased compared with WT mice at 15-24-week-old (Figure 3B). We demonstrated that the IL-6 mRNA levels were higher in intestinal tumors of Apc^Min/+^ mice than in normal intestinal epithelium of WT mice by qPCR (Figure 3C). Previous study reported MC38 cells and CT26 cells all can secret IL-6(Li et al., 2018). Therefore, tumor cells can be the source of IL-6. In vitro, we used protein microarrays to detect inflammatory factors in the supernatant of mouse colorectal carcinoma MC38 cell lines and the results showed that IL-6 expression was higher in the supernatant after cultured with MC-38 cells (Figure 3D, E). So, tumor cells can secrete IL-6.

**Figure 3.**
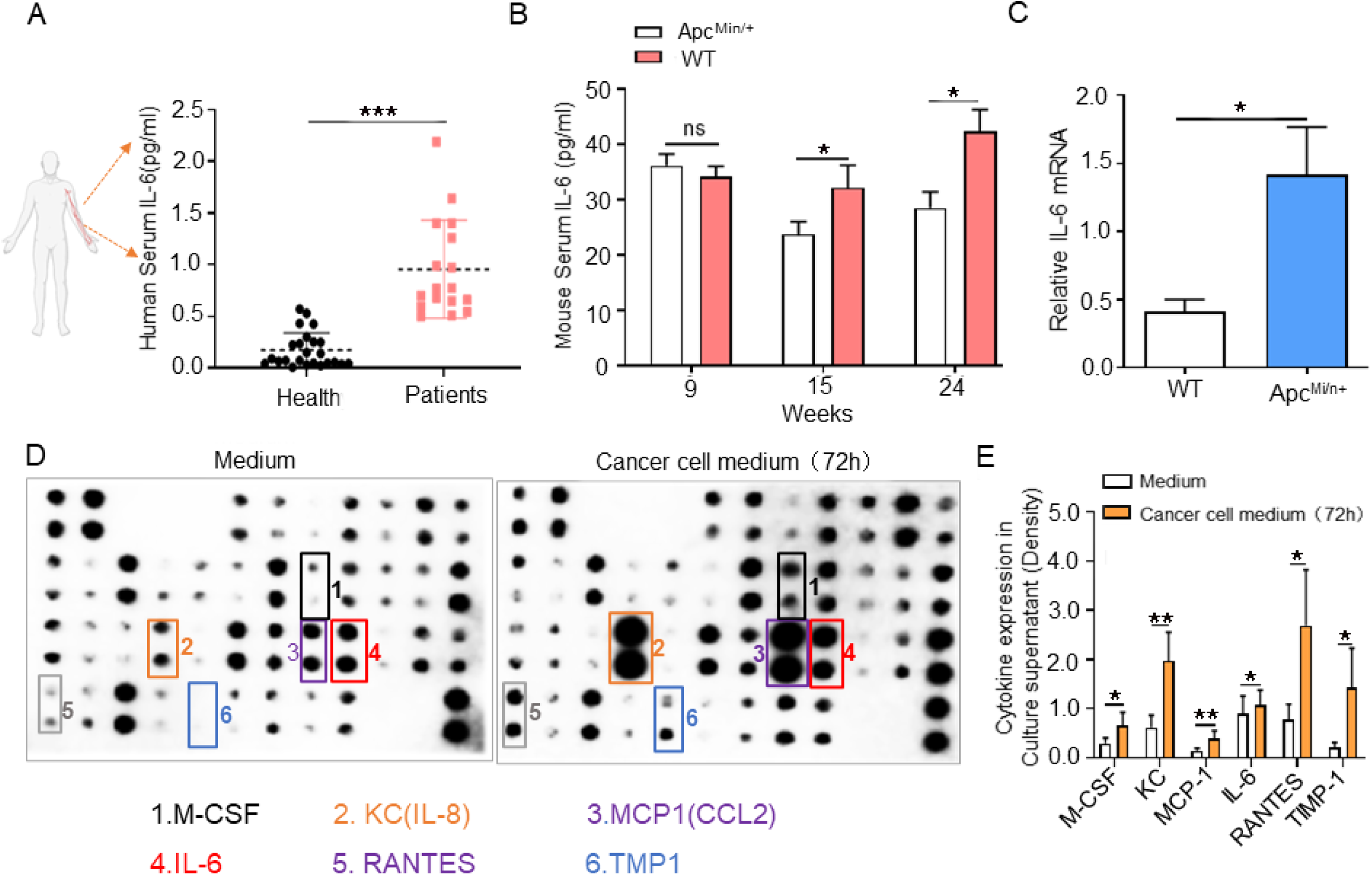
Tumor cells can secrete interleukin 6. (A)Serum interleukin 6 (IL-6) levels in normal individuals and patients with colorectal cancer aged 30-50 years detected by enzyme-linked immunosorbent assay (\*\*\**P<*0.001; data are shown as the means ± SD; n = 26 per group). (B) The IL-6 levels in serum in WT mice and Apc^Min/+^ mice were detected at 9-, 15-, and 24 weeks old by enzyme-linked immunosorbent assay (\**P<* 0.05; data are shown as the means ± SD; n = 5 per group). (C) IL-6 mRNA expression was validated in normal intestinal epithelium isolated from Apc^Min/+^mice versus that in intestinal tumors isolated from WT mice by qPCR (\**P<* 0.05; data are shown as the means ± SD; n =4 per group). (D) Cytokine array detects inflammatory cytokines in fresh untreated medium (-MC38 cells) and cultured MC-38 cells (+MC38 cells); arrows indicate the significantly increased cytokines. (E) The relative quantification of the significantly upregulated cytokine to positive quality control density ratio by ImageJ software. The positive quality control density was determined for normalization purposes (\*\**P<* 0.01, \**P<* 0.05; data are shown as the means ± SD).

### 3. 4 Tumor-derived IL-6 can upregulate MMP12 in macrophage

Because some previous studies reported that IL-6 may not directly lead to muscle loss in CAC(Carson and Baltgalvis, 2010a; Franckhauser et al., 2008). IL-6 in the tumor microenvironment may be an important determinant of alternative macrophage activation and induce macrophage M2 polarization and M2 macrophages can produce MMP12(Suzuki et al., 2017; Wang et al., 2018). Taken together, we speculated whether tumors regulate macrophage MMP12 by secreting IL-6 to affect muscle loss. To uncover the underlying mechanism communication between tumor-derived IL-6 and macrophages, we performed cell experiments in vitro (as described in the Materials). Mouse macrophage RAW264.7 cells co-cultured with mouse colorectal cancer MC38 cells (CT 26 cells) for 72hours to detect macrophage MMP12 by western blotting (Figure 4A) and the results confirmed that RAW264.7 cells exhibited increased MMP12 expression as the number of MC38 cells increased, with RAW264.7 cells cultured alone as the negative control group (Figure 4B, C). Similar trends were observed in CT26 cells (Figure 4D, E). We further treated RAW264.7 cells with IL-6 in different methods. RAW264.7 cells were seeded into 6-well plates and treated with increasing doses of IL-6 (0, 2, 5 10,30ng/ mL) for 72h. Next, RAW264.7 cells were treated continuously with IL-6 (30 ng/ml) for 0, 3, 6, and 9hours. Cells incubated with fresh media were used as the untreated negative controls (Figure 4F). We found that within a certain concentration range (<30ng/ml), as the IL-6 dose increased, the expression of MMP 12 in RAW264.7 cells also increased when treated with IL-6 (Figure 4G, H). Next, the expression of MMP12 in RAW264.7 increased as the stimulation time prolonged when the RAW264.7 cells treated with IL-6(30ng/ml) (Figure 4I, J).

**Figure 4.**
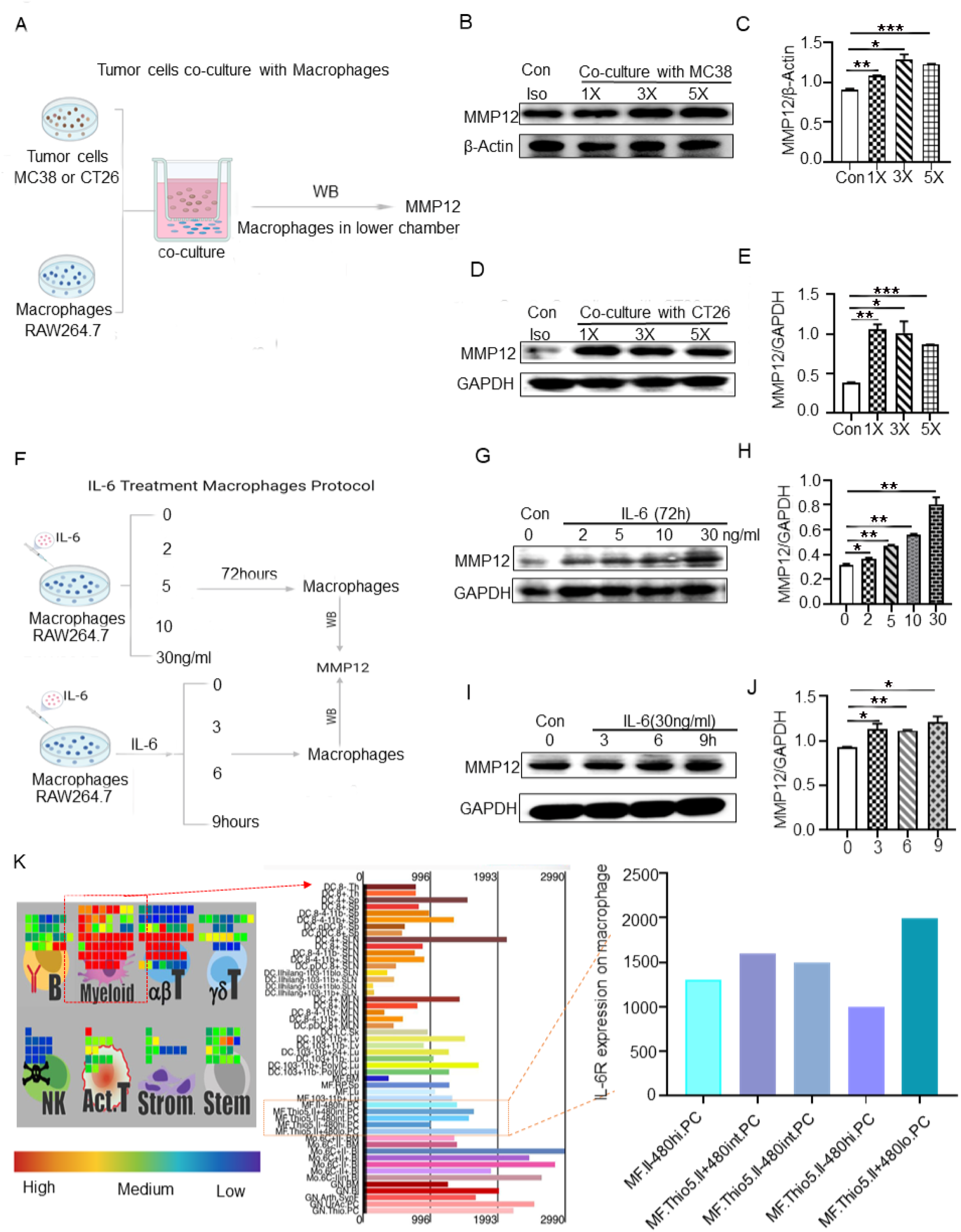
Tumor-derived IL-6 can upregulate MMP12 in macrophages. (A)Schematic diagram of tumor cells (MC38/CT26 cell lines) coculture with macrophage cells (RAW264.7 cell lines). All quantifications use image J software grayscale statistics. (B, C) Representative western blots showing the secreted MMP12 protein levels from RAW264.7 cell lines (1-2×10^5^) cultured alone or cocultured with MC38 cell lines (control, 1×10^4^, 3×10^4^, 5×10^4^). β-Actin was used as the internal control for normalization purposes. (D, E) Representative western blots showing the secreted MMP12 protein levels from RAW264.7 cell lines (1-2×10^5^) cultured alone or cocultured with CT26 cell lines (control, 1×10^4^, 3×10^4^, 5×10^4^). GAPDH was used as the internal control for normalization purposes. (F) Schematic diagram of IL-6 treated macrophages. RAW264.7 cells incubated with fresh media were served as untreated negative controls. Using western bloting to detect MMP12 in RAW 264.7 cells and GAPDH was used as the internal control. (G, H) RAW264.7 cells (1-2×10^5^) were seeded into 6-well plates and treated with increasing doses of IL-6 (0, 2, 5 10,30ng/ mL) for 72hours. (I, J) RAW264.7 cells (1-2×10^5^) were treated continuously with IL-6 (30 ng/ml) for 0, 3, 6, and 9hours. (K) Immune gene data proved that IL-6 receptor is expressed on myeloid cells and the red box represents F480^+^ macrophages. The colored bars refer to the expression level of IL-6 receptors on macrophages. Red represents high expression of IL-6 receptors, and green represents low expression of IL-6 receptors.

Meanwhile, immune gene data proved that IL-6 receptor (IL-6R) is highly expressed on myeloid cells, including F480^+^ macrophages (Figure 4K). The previous studies proved that IL-6 can be derived from MC38 and CT26 tumor cells (Li et al., 2018). Taken together, these findings suggest that tumor-derived IL-6 can stimulate macrophages and up-regulate MMP12 in macrophages.

### 3.5 MMP12 can degrade insulin and insulin-like growth factor-1

The present results have proved that tumor-derived IL-6 can up-regulate MMP12 in macrophages. Knockout of MMP12 can reduce muscle loss in Apc^Min/+^ mice. Recently, it has been demonstrated that insulin and insulin-like growth factor 1(IGF-1) have complex anabolic effects and are important regulators of muscle remodeling that can mediate muscle atrophy(Baker Rogers et al., 2020; Dev et al., 2019; Han et al., 2019; Masi and Patel, 2020; Takayama, 2019). Moreover, Jung-Ting Lee proposed that MMP12 expression significantly promoted insulin resistance and that insulin may be regulated by resident macrophages(Lee et al., 2014a). To understand the molecular mechanism underlying muscle loss by macrophage MMP12, we further examined the relationship between MMP12 and insulin (IGF-1) which affects muscle loss. Firstly, after the labeled insulin polypeptide is incubated with serum, the absorbance increases (λ=488nm) (Figure 5A). Because IGF-1 is similar in structure to insulin, furthermore, we verified the relationship between fluorescently labeled insulin (IGF-1) and MMP12, and measured its fluorescence intensity and characteristic peak changes (Figure 5B). We found that when the dose of insulin fluorescent peptide is constant, the more MMP12 protein, the stronger the fluorescence intensity. Similar Tendencies were observed in the IGF-1(Figure 5C). The qualitative results of electrospray ionization mass spectrometry showed that the characteristic peak of insulin fluorescent peptide (m/z = 436.99) disappeared after incubation with MMP12 protein (Figure 5-figure Supplement 2A). When the IGF-1 polypeptide was incubated with MMP12, its characteristic peak (m/z = 436.68) disappeared (Figure 5-figure Supplement 2B). Taken together, MMP12 can indeed degrade insulin and IGF-1. It seems that the degradation of MMP 12 to IGF-1 is stronger than that of insulin.

**Figure 5.**
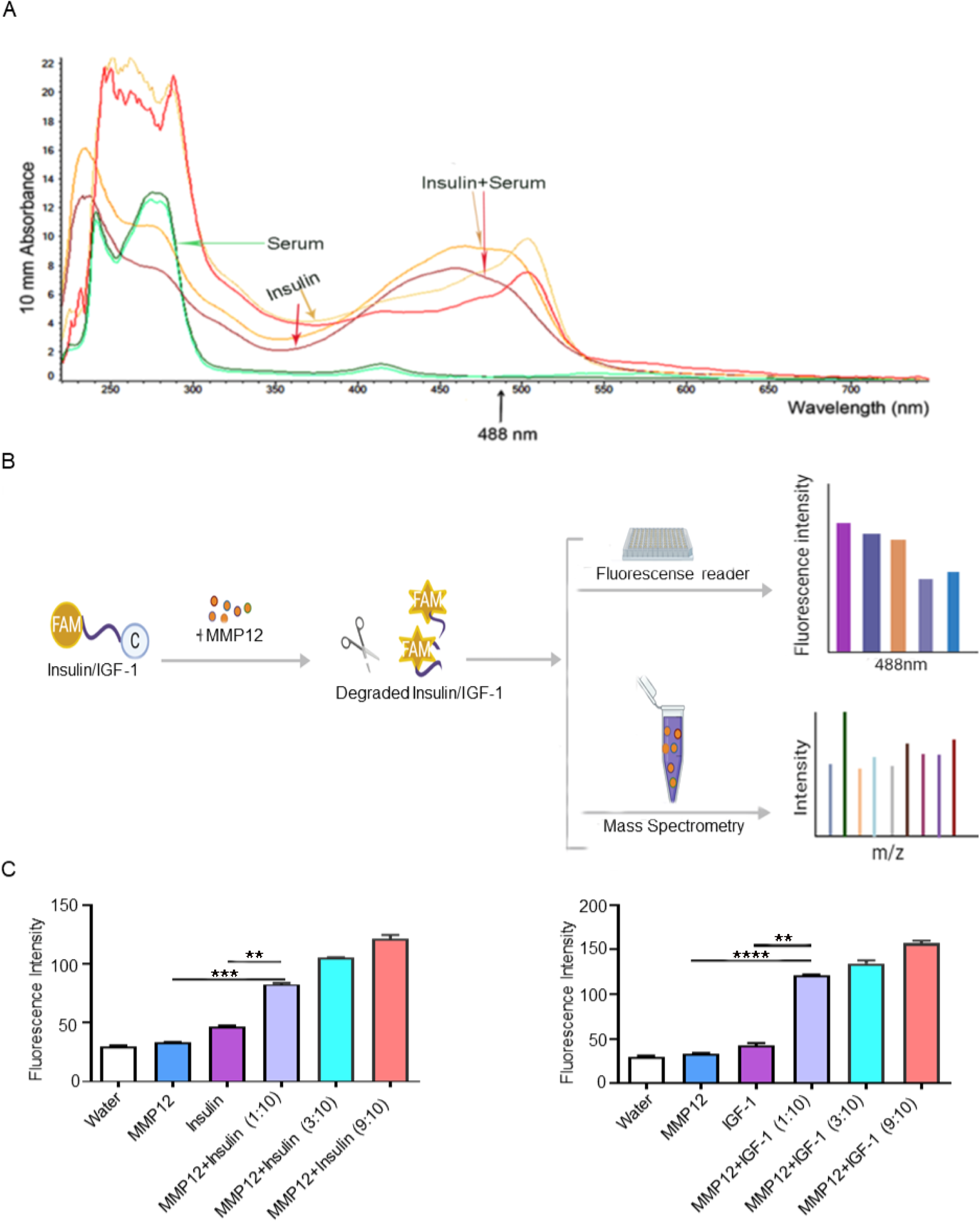
MMP12 can degrade insulin and insulin-like growth factor-1. (A) Representative picture of the peak shift after the insulin polypeptide interacts with serum. (A) The synthetic insulin (or insulin-like growth factor-1) peptide was labeled with FAM and DABCLY as shown; if the insulin (or insulin-like growth factor-1) peptide was degraded, the FAM signal was detected. This is based on fluorescence resonance energy transfer (FRET); Detection of characteristic peaks of insulin alone and the mixture of insulin (or insulin-like growth factor-1) and MMP12 by ionization mass spectrometry. (C)The coexistence of MMP12 and insulin (or insulin-like growth factor-1) peptide led to a fluorescence signal and appeared dose-dependent (\*\*\**P<* 0.001, \*\**P<* 0.01; data are shown as the means ± SD).

### 3.6 MMP12 inhibitor can rescue weight loss of Apc^Min/+^ mice

It is reported that insulin and insulin-like growth factor 1 can indeed affect the muscle loss caused by cachexia and exacerbate weight loss(Baker Rogers et al., 2020; Dev et al., 2019; Han et al., 2019; Masi and Patel, 2020; Takayama, 2019). Therefore, we are concerned about whether the inhibition of MMP12 that degrades insulin and insulin-like growth factor 1 affects the weight change of cachexia mice. To investigate the effect of inhibiting MMP12 on colorectal cancer Apc^Min/+^ mice, we combined the MMP12 inhibitor (MMP408) and a classic clinical anti-colon cancer drug (5-FU) (Figure 6A). After 2 weeks of administration in Apc^Min/+^ mice, at 17-week-old, the results showed that the weight loss in the MMP12 inhibitor group (+MMP408) only accounted for 5% of the basal body weight and was only one third of that of the control normal saline group (+Control). There was a significant difference between the two groups. In the MMP12 inhibitor and anticancer drug combination group (+MMP408+5-FU), the weight change was decreased by approximately 8% of the basal body weight and was half that of the normal saline group (+Control). Unfortunately, there were no changes in body weight in the anticancer drug combination group (+MMP408+5-FU), when compared with the MMP12 inhibitor group alone (+ MMP408) (Figure 6B). In summary, the above experiments proved that specifically inhibiting MMP12 at the CAC stage of weight loss in Apc^Min/+^ mice can reduce weight loss.

**Figure 6.**
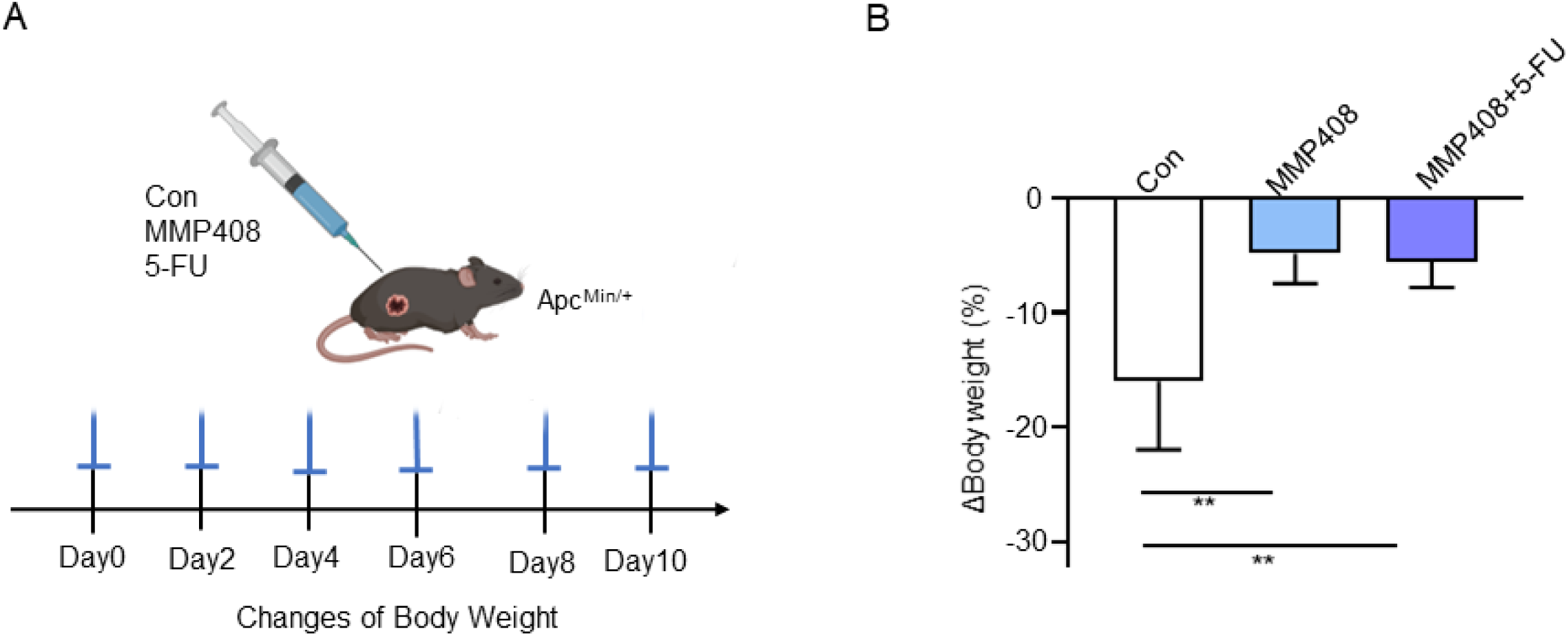
Inhibiting MMP12 in Apc^Min/+^ mice reduces weight loss. (A)Schematic diagram of the administration process of 17-week Apc^Min/+^ mice. The drug was given every two days (MMP408-5mg/kg, 5-FU-30mg/kg). The saline group was used as a control. (B) Percentage of weight gain compared to the basal weight after administration of drugs in Apc^Min/+^ mice (\*\**P<* 0.01, data are shown as the means ± SD; n = 5 per group).

## 4 Discussion

Researches on weight loss have received more and more attention in many fields, such as diabetes, abnormal thyroid metabolism, weight control, etc, among which, muscle loss caused by malnutrition similar to cancer cachexia (CAC) is more worrying. More than four-fifths of patients with CAC die from extreme loss of body weight and skeletal muscle. Our study suggested that MMP12 plays a new role in controlling weight and muscle loss and inhibiting MMP12 can reverse the body weight reduction with CAC. In details, the major findings of our study are as follows: (1) MMP12 promotes weight loss and accelerates the deterioration of CAC. The loss of weight and muscle induced by CAC was reduced in Apc^Min/+^ by MMP12 knockout. (2) In vivo, MMP12-positive immunostaining was found in muscle of human and mice. MMP12 was co-labeled with macrophages in the muscle tissue in situ. Importantly, MMP12 positive staining was substantially increased in the muscle and peritoneal macrophages from Apc^Min/+^ mice compared with those from wild type (WT) mice. (3) Clinically, serum interleukin 6 (IL-6) increased in cancer patients. A similar increasing trend was found in serum and tumor tissues of Apc^Min/+^ mice compared to those from WT mice. Crucially, IL-6 has been shown to be directly secreted by MC38 tumor cells. (4) In vitro, we proved that tumor cells have a positive relationship with MMP12 secreted by macrophages. At the cellular level, we found for the first time that macrophages can be stimulated and regulated by IL-6, and the level of MMP12 in macrophages was up-regulated. (5) Underlying mechanism, MMP12 can degrade insulin and insulin-like growth factor 1. In our study, the degradation effect of MMP12 on insulin-like growth factor 1 was proved for the first time, and it was found that the degradation effect of MMP12 was stronger than that of insulin. (6) Inhibiting MMP12 prevent weight loss in Apc^Min/+^ mice at CAC stage. In summary, the present study uncovered a novel mechanism that MMP12 promotes weight and muscle loss. The crosstalk between tumor cells and macrophages is that MMP12 is upregulated by tumor cell-derived IL-6 and MMP12 can degrade insulin and insulin-like growth factor 1, affecting glycolipid metabolism, resulting in weight loss (Figure 7).

**Figure 7.**
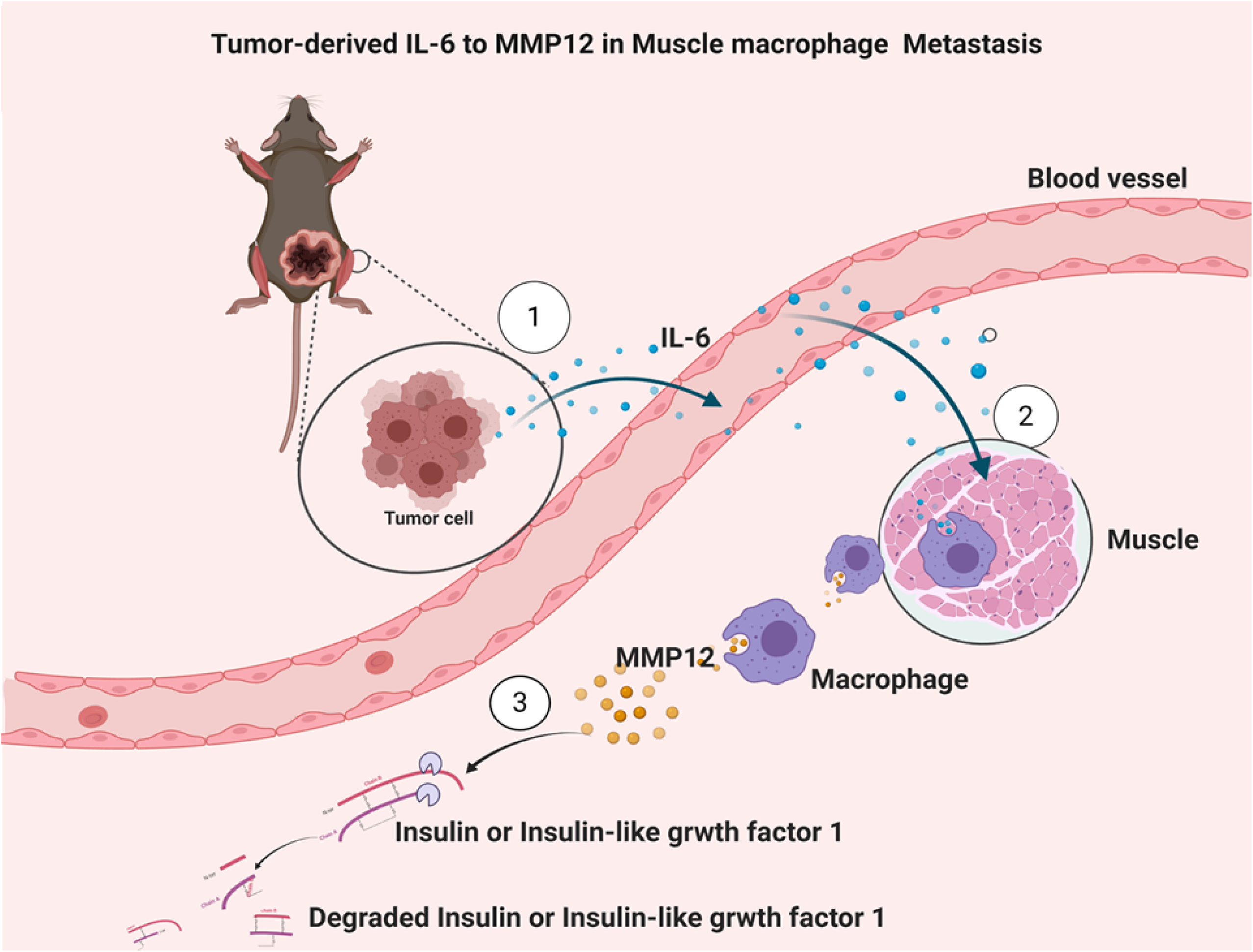
The role of MMP12 in the crosstalk between tumor and muscular macrophage in cancer cachexia.

Weight and muscle loss induced by CAC is main cause of death in cancer patients worldwide. The clinical definition of CAC by Fearon criteria includes the following characteristics: weight loss > 5% or weight loss > 2% and a BMI < 20 kg/m^2^ or sarcopenia(van der Werf et al., 2018). In our study, 15-24-weeks old Apc^Min/+^ mice with weight loss > 15%, as well as muscle (gastrocnemius and soleus) loss, and certain other symptoms such as anemia, were considered as the characteristics of CAC. Our data exhibited that muscle weight and muscle cross-sectional area increased in Apc^Min/+^; MMP12^-/-^ mice compared with Apc^Min/+^ mice, which indicating that knocking out MMP12 may suppress the decrease of skeletal muscle.

As a kind of matrix metalloproteinase family, MMP12 is also called macrophage elastase. It was previously reported that MMP12 can decompose various extracellular matrix components and vascular components, and MMP12 is involved in tumor cell invasion and metastasis. In 2014, Lee Jung-Ting had explored the role of MMP12 in white adipose tissue expansion on high-fat feeding(Jung-Ting et al., 2014), while the function of MMP12 under the lack of nutrition yielded has not been studied. We firstly tried to build a model with tumor-bearing environment lacking nutrition in Apc^Min/+^ mice hybridized with MMP12^-/-^ mice. Unexpectedly, under the condition of tumor-bearing mice, knocking out MMP12 caused a decrease in muscle loss, but not affect white adipose tissue.

A mRNA analysis of the data got from The Cancer Genome Atlas (TCGA) for GTEx, Illumina, BioGPS and SAGE of MMP12 gene in normal human tissues (Figure 1-figure Supplement 3C). In our study, we focus on the liver, muscle and fat tissues. We weighed these mice tissues at 24-weeks old, and performed histological evaluation using H&E staining. The liver weight has difference between Apc^Min/+^ mice and Apc^Min/+^; MMP12^-/-^ mice (Figure 1-figure Supplement 4 A), however, H&E staining showed that knocking out MMP12 had no more histology effect on the WT mice and Apc^Min/+^ mice (Figure 1-figure Supplement 4B). Similarly, knocking out MMP12 in Apc^Min/+^ mice did not cause changes in white fat, even though knocking out MMP12 in the wild background can indeed cause white fat increase and expansion, which in agreement with Lee Jung-Ting, who showed in 2014 that knocking out MMP12 can increase fat expansion when performing high-fat feeding in WT mice(Jung-Ting et al., 2014). Notably, the ratio of brown adipose tissue-to-body weight decreased in Apc^Min/+^ mice compared with Apc^Min/+^; MMP12^-/-^ mice (Figure 1-figure Supplement 4C), and the area of brown adipose tissue in MMP12^-/-^ mice expansion, and serving the Apc^Min/+^ mice as the control group, the same tendency showed in Apc^Min/+^; MMP12^-/-^ mice (Figure 1-figure Supplement 4D), suggesting there may be a tendency to convert to white adipose tissue. However, the reasons have not been extremely explored in the current research. We hypothesized that MMP12 may play an important role in the conversion of brown adipose tissue and white adipose tissue. and it may be associated with a CAC energy consumption and even as useful for researcher focusing on weight loss drug. MMP12 is mainly derived from macrophages. In view of the fact that I have found that knocking out MMP12 can increase muscle weight and cross-sectional area, we conducted related experiments on whether MMP12 is expressed on muscles. Studies have shown that MMP12 is existed and co-labeling with macrophages in our study. Notably, MMP12 levels increase in muscle tissues and peritoneal macrophages not in serum.

IL-6 is mainly secreted by a variety of immune cells and is also highly expressed in a variety of cancer cells(Mauer et al., 2015). Our in vitro studies proved that MC38 cell lines can secrete IL-6. Our animal experiments in vivo confirmed that the serum IL-6 of Apc^Min/+^ mice was also higher than that of WT mice at 15 weeks and 24 weeks, which is consistent with the previous study(Baltgalvis et al., 2008). IL-6 mRNA levels in intestinal tumors are increased compared with normal intestinal epithelial tissue, which echoes our data with the increased serum IL-6 in clinical tumor patients(Nikiteas et al., 2005). In short, maybe, the increased IL-6 in the tumor then circulates into the blood. Of course, the cytokines secreted by MC38 cells also include monocyte chemoattractant protein1(MCP1) and keratinocyte-derived chemokine (KC) which can recruit macrophages(Barcelos et al., 2004; Engin, 2017; Wang et al., 2011). But unfortunately, our experiments have shown that only increased at mRNA levels of intestinal tumors in Apc^Min/+^ mice at CAC stage but serum MCP1did not change at the CAC stage in Apc^Min/+^ mice (Figure 3-figure Supplement 5A, B). There was no difference in serum KC and KC mRNA levels in late tumors in mice (Figure 3-figure Supplement 5C, D). Similarly, clinically, serum KC did not differ between normal healthy individuals and colorectal cancer patients (Figure 3-figure Supplement 5E).

In summary, we suspected that there may be a possibility that MCP1 and KC can also cooperate with IL-6 to recruit macrophages, thereby activating alternative macrophages to polarize M2 macrophages, resulting in MMP12 secretion. The above results suggested that in the period of CAC, the key is that IL-6 secreted by tumor cells plays a major role(Baltgalvis et al., 2008). In Table 2.

**Table 2.**
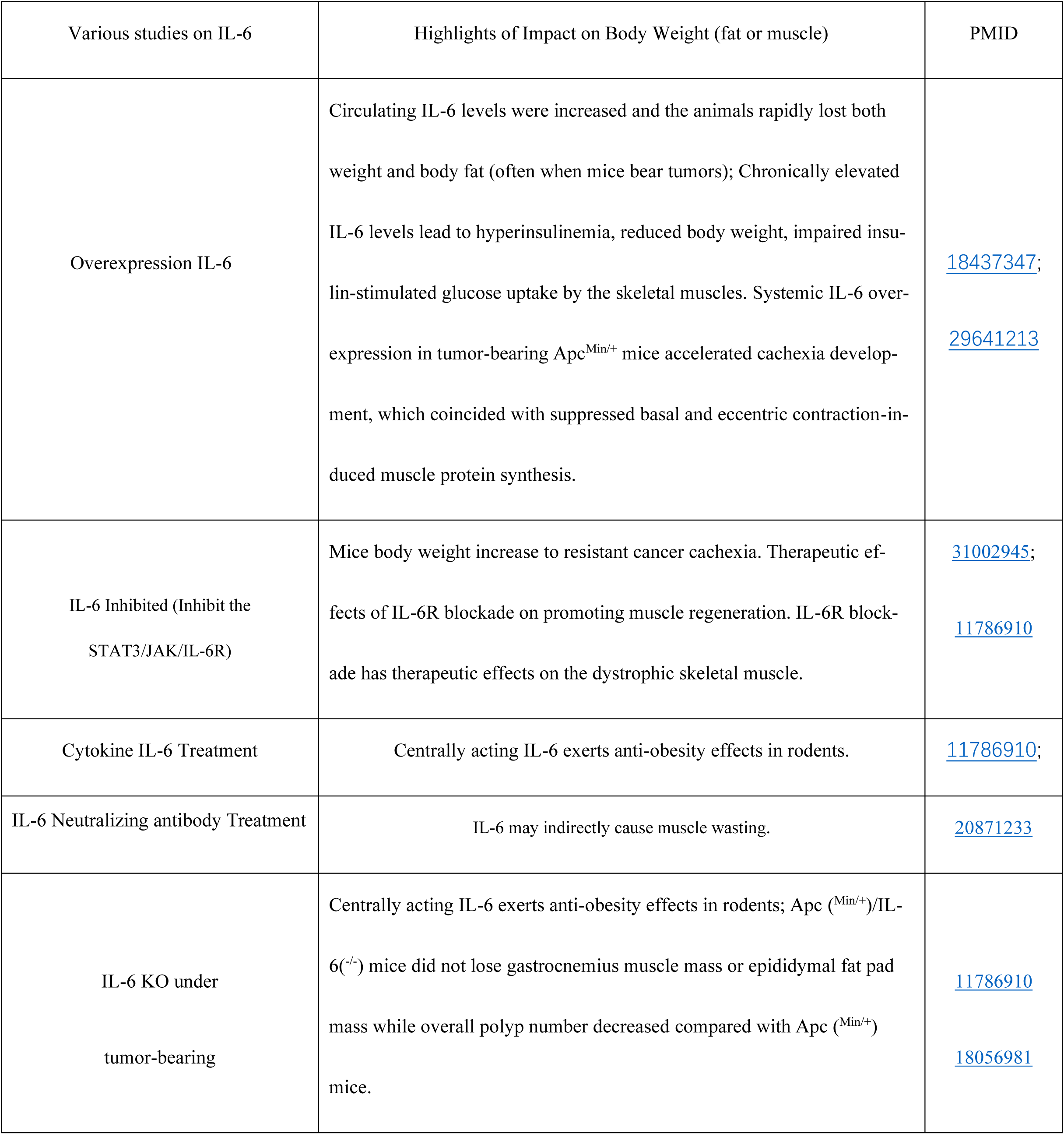
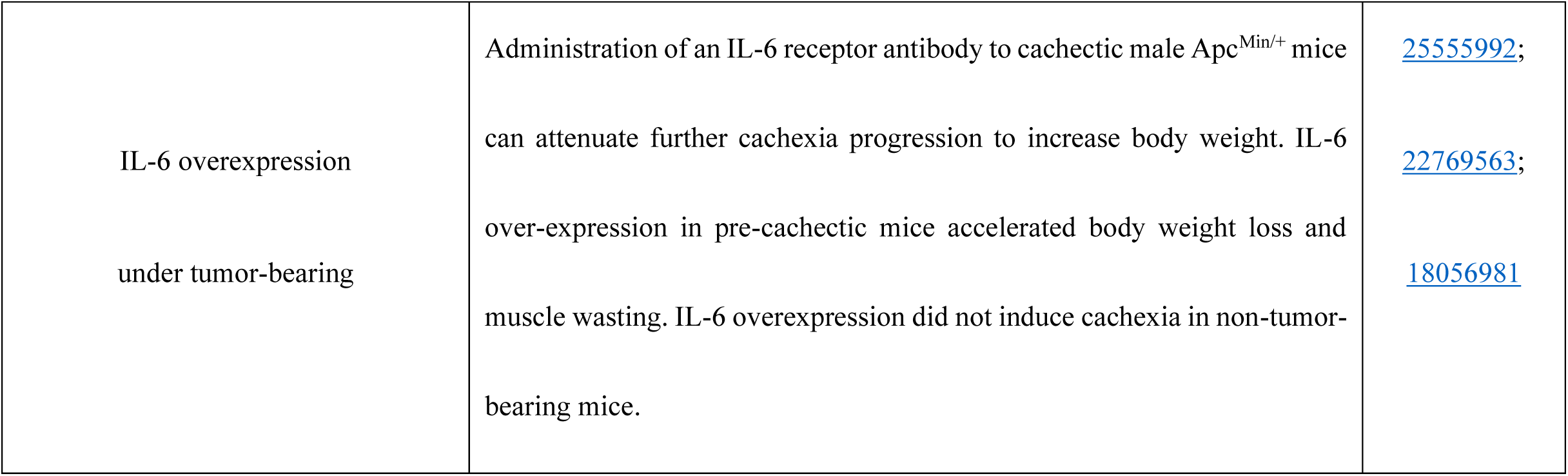
The effect of IL-6 on body weight.

| S Table 2 |  |  |
| --- | --- | --- |
| Various studies on IL-6 | Highlights of Impact on Body Weight (fat or muscle) | PMID |
| Overexpression IL-6 | Circulating IL-6 levels were increased and the animals rapidly lost both weight and body fat (often when mice bear tumors); Chronically elevated IL-6 levels lead to hyperinsulinemia, reduced body weight, impaired insulin-stimulated glucose uptake by the skeletal muscles. Systemic IL-6 overexpression in tumor-bearing Apc<sup>Min/+</sup> mice accelerated cachexia development, which coincided with suppressed basal and eccentric contraction-induced muscle protein synthesis. | 18437347;<br>29641213 |
| IL-6 Inhibited (Inhibit the STAT3/JAK/IL-6R) | Mice body weight increase to resistant cancer cachexia. Therapeutic effects of IL-6R blockade on promoting muscle regeneration. IL-6R blockade has therapeutic effects on the dystrophic skeletal muscle. | 31002945;<br>11786910 |
| Cytokine IL-6 Treatment | Centrally acting IL-6 exerts anti-obesity effects in rodents. | 11786910; |
| IL-6 Neutralizing antibody Treatment | IL-6 may indirectly cause muscle wasting. | 20871233 |
| IL-6 KO under tumor-bearing | Centrally acting IL-6 exerts anti-obesity effects in rodents; Apc<sup>(Min/+)</sup>/IL-6<sup>(-/-)</sup> mice did not lose gastrocnemius muscle mass or epididymal fat pad mass while overall polyp number decreased compared with Apc<sup>(Min/+)</sup> mice. | 11786910<br>18056981 |
| IL-6 overexpression<br>under tumor-bearing | Administration of an IL-6 receptor antibody to cachectic male Apc<sup>Min/+</sup> mice can attenuate further cachexia progression to increase body weight. IL-6 over-expression in pre-cachectic mice accelerated body weight loss and muscle wasting. IL-6 overexpression did not induce cachexia in non-tumor-bearing mice. | 25555992;<br>22769563;<br>18056981 |

**Table 3:** Abbreviations.

| **Abbreviations** | **Full name** |
| --- | --- |
| MMP12 | Matrix Metalloproteinases 12 |
| CRC | Colorectal Cancer |
| CAC | Cancer Cachexia |
| Apc<sup>Min/+</sup>; MMP12<sup>-/-</sup> | Apc<sup>Min/+</sup>; MMP12<sup>-/-</sup> mice |
| WAT | White Adipose Tissue |
| BAT | Brown Adipose Tissue |
| HRP | Horseradish Peroxides |
| H&E | Hematoxylin & Eosin |
| qPCR | Quantitative Polymerase Chain Reaction |
| IHC | Immunohistochemistry |
| IF | Immunofluorescence |
| PBS | Phosphate Buffered Saline |
| MCP1(CCL2) | Monocyte Chemoattractant Protein 1 |
| KC(CXCL1) | Keratinocyte-derived Chemokine |
| RANTES | Regulated upon Activation, Normal T Cell Expressed and Presumably Secreted |
| IL-6 | Interleukin 6 |
| IGF-1 | Insulin-like Growth Factor 1 |

we summarized the research on IL-6 on body weight and muscle loss, which mostly demonstrated that IL-6 may have indirect effect on body weight and muscle. Especially, Kristen A Baltgalvis pointed out knocking out IL-6 can reduce muscle consumption in Apc^Min/+^ mice(Baltgalvis et al., 2008).

The crosstalk between tumors and inflammatory factors is well known(Talbert et al., 2018b; Zhang et al., 2008). We use co-culture experiments in MC38 cell lines and CT26 with RAW264.7 cell lines to demonstrate that macrophage MMP12 increase treatment with IL-6. We speculated that this is caused by the IL-6 secreted by tumor cells, and we firstly proved that IL-6 can directly upregulate macrophage MMP12. However, we did not determine whether the IL-6 produced by macrophages acts on macrophages themselves. Similarly, we have not explored whether other cytokines secreted by tumors or macrophages themselves play a synergistic or indirect role along with IL-6.

In present study, we used fluorescence intensity measurement and ionization mass spectrometry methods to prove that insulin and IGF-1 can indeed be degraded by MMP12, but the specific sites and amino acids where insulin and IGF-1 were broken have not been further proved. At the same time, we could not prove to completely rule out whether MMP12 itself generates characteristic peaks from fracture (Figure 5-figure Supplement 2A, B).

MMP12, as a macrophage matrix metalloproteinase, has been repeatedly reported to degrade insulin and affect insulin sensitivity. We verified MMP12 can degrade insulin or IGF-1 in vitro, which is consistent with the previous study(Kettner et al.). As the two key hormones in tumor microenvironment, insulin resistance is correlated with in insufficient insulin, lack of insulin receptor, or decreased insulin sensitivity, which will reduce the uptake of glucose in organs, which suggests that MMP12 is closely related to glycolipid metabolism, leading to the loss of skeletal muscle and adipose tissue(Baker Rogers et al., 2020; Dev et al., 2019; Han et al., 2019; Masi and Patel, 2020; Takayama, 2019). The insulin kits and insulin tolerance test and oral glucose tolerance test results showed that the knocking out MMP12 in Apc^Min/+^ mice may reduce insulin levels or increase insulin sensitivity, reversing insulin resistance (Figure 5-figure Supplement 6A-H), but is not related to basic function of islets according to H&E staining and IHC staining (Figure 5-figure Supplement 4E-H). We tested four blood lipid levels with the kit, and the results showed that when Apc^Min/+^ mice were knocked out MMP12, total triglycerides decreased in the early and middle stages, but high density lipoprotein cholesterol increased in all age groups, total cholesterol and low density lipoprotein cholesterol not changed across the 4 groups (Figure 5-figure Supplement 6I-L). So, does IL-6 regulating MMP12 help restore muscle loss caused by cachexia? Clinical inhibiting IL-6 may reduce CAC patients with weight loss. However, long-term treatments with high-dose IL-6 may cause additional side effects, such as exacerbating CAC resulting in more muscle loss(Wada et al., 2017). After all, IL-6 acts on muscles indirectly. Surprisingly, MMP12, as the downstream of IL-6, can significantly suppress weight loss when being specifically inhibited in mice, although the effect is not more obvious after combined treatment with the classic colorectal cancer chemotherapy drug 5-FU. Clinically, suppressing MMP12 may reduce the possibility of insulin degradation, suggesting that our findings may as a method to treat directly glucose deficiency, CAC and complications of CAC. It will not only provide a new direction for reducing the blood glucose and blood lipid levels of cancer patients but also bring new research ideas for the clinical treatment of diabetes caused by insulin deficiency.

In summary, our results identified that knocking out MMP12 in Apc^Min/+^ mice significantly reduced muscle loss caused by CAC. We determined a positive correlation with between tumor-derived IL-6 and macrophage MMP12 in colorectal cancer. MMP12 can degrade insulin and IGF-1, reversing the insulin resistance in CAC to regulate tumor glycolipid metabolism. Therefore, MMP12 is a double-edged sword for tumor microenvironment, but, inhibiting MMP12 may represent a new potential targeting for the treatment of clinical patients with weight loss.

## Authors’ Contributions

Conceptualization: Jiangchao Li, Lijing Wang, Lingbi Jiang

Methodology: Jiangchao Li, Lijing Wang, Lingbi Jiang,

Software: Lingbi Jiang

Validation: Lingbi Jiang, Mingming Yang, Ting Niu

Formal analysis: Jiangchao Li, Lingbi Jiang

Investigation: Jiangchao Li, Lingbi Jiang,

Resources: Zhengyang Li, Lili Wei, Haobin Li, and Ting Niu, Mingzhe Huang, Pinzhu Huang

Data curation: Lingbi Jiang, Jiangchao Li

Writing – original draft preparation: Lingbi Jiang, Jiangchao Li,

Writing – review & editing: Visualization; Jiangchao Li, Lijing Wang, Yan Mei, Rongxin Zhang, Dehuan

Xie, Yan Mei

Supervision; Jiangchao Li, Lijing Wang,

Project administration; Lingbi Jiang and Shihui He, Zhengyang Li, Xiaodong He

Funding acquisition:Jiangchao Li, Lijing Wang, Rongxin Zhang. This work was supported by grants from the National Natural Science Foundation of China (Grant ID: 81773118 to Jiangchao Li, 31771578 to Lijing Wang and 81872320 to Rongxin Zhang). The funding source involves research design, data collection and interpretation, and now it is decided to submit the work for publication.

## Acknowledgments

We thank technician Dr. Hao Chen for their help in clinical sample collection. We appreciate that Prof. Ming Li gave us advices for this study. We would also like to thank Jingzhou Xie, Lixun Huang, Yongjia Zheng, Yiting Zhang, Junwei Ye, Qianhui Ma, Jiena Liu and Xiaoyang Chen for their help with animal experiments.

## Ethics Statement for Human Subjects Research or Animal Experimentation

All mouse experimental protocols were approved by the animal experimental ethics committee of Guangdong Pharmaceutical University. The animal ethics approval number was gdpulac2019019. All tests were carried out with the approval of the Guangdong Medical Laboratory Animal Center, Guangzhou, China. All experiments for clinical patients in this study were obtained by the approval of the Guangzhou Human Research Ethics Committee, Provincial First Affiliated Hospital of Guangdong Pharmaceutical University, China. The clinical ethics approval number was EC-AF-019.

## Supplementary data

**Figure S1.**
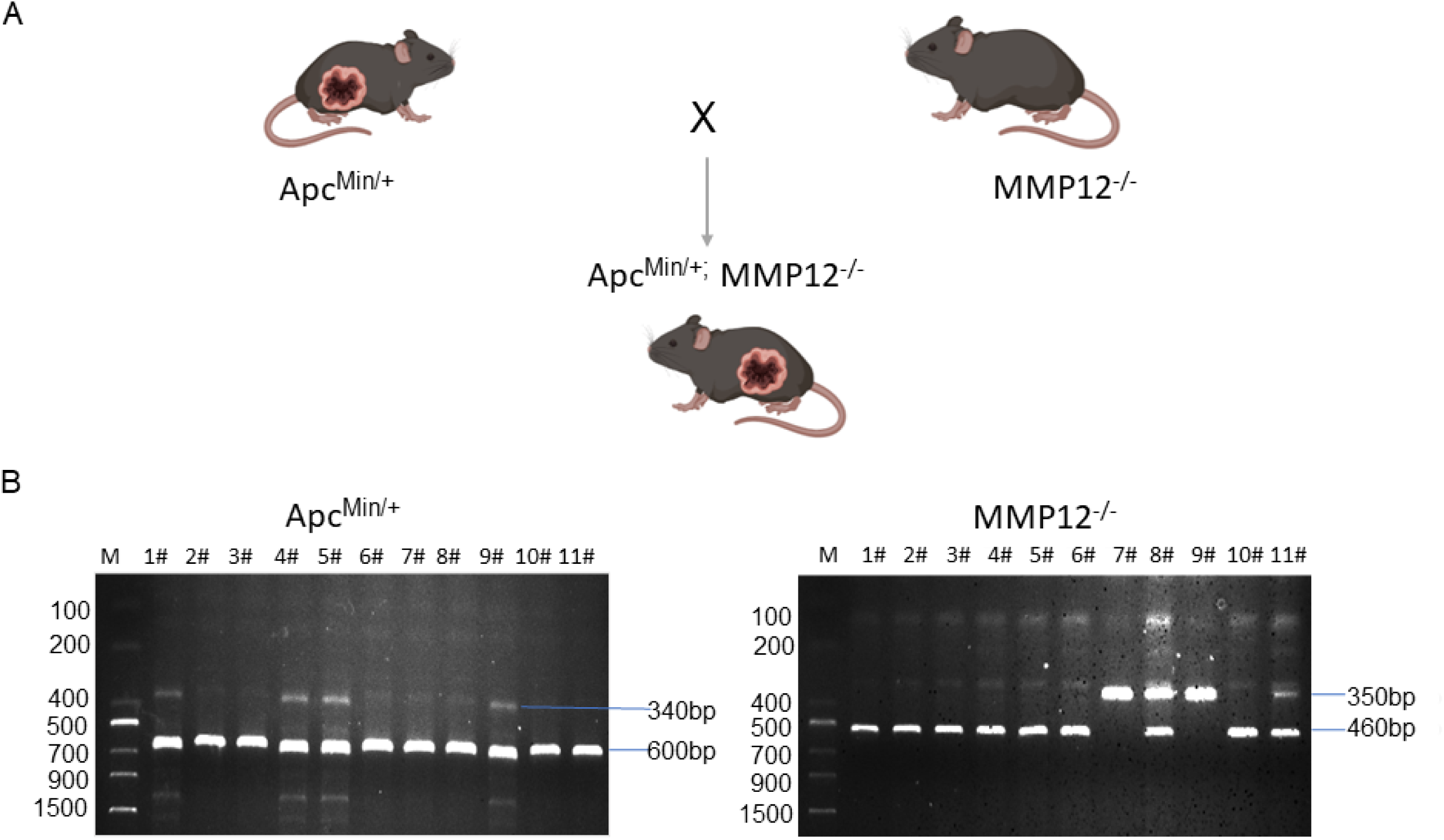
Mouse crossbreeding and genotype identification. A. Schematic of the crossbreeding of Apc^Min/+^ mice with MMP12^-/-^ mice to obtain Apc^Min/+^; MMP12^-/-^ mice. B. The APC gene mutant PCR product size was 340 bp, and the PCR product size of wild-type (WT) mice was 600 bp. The MMP12 knockout (mutation) PCR product size was 460 bp, and the WT (WT) PCR product size was 350 bp. In detail, Apc^Min/+^:1#, 4#, 5#, 9#; MMP12^-/-^:1#, 2#, 3#, 4#, 5#, 6#, 10#, 11#; Apc^Min/+^; MMP12^-/-^:1#, 4#, 5#.

**Figure S2.**
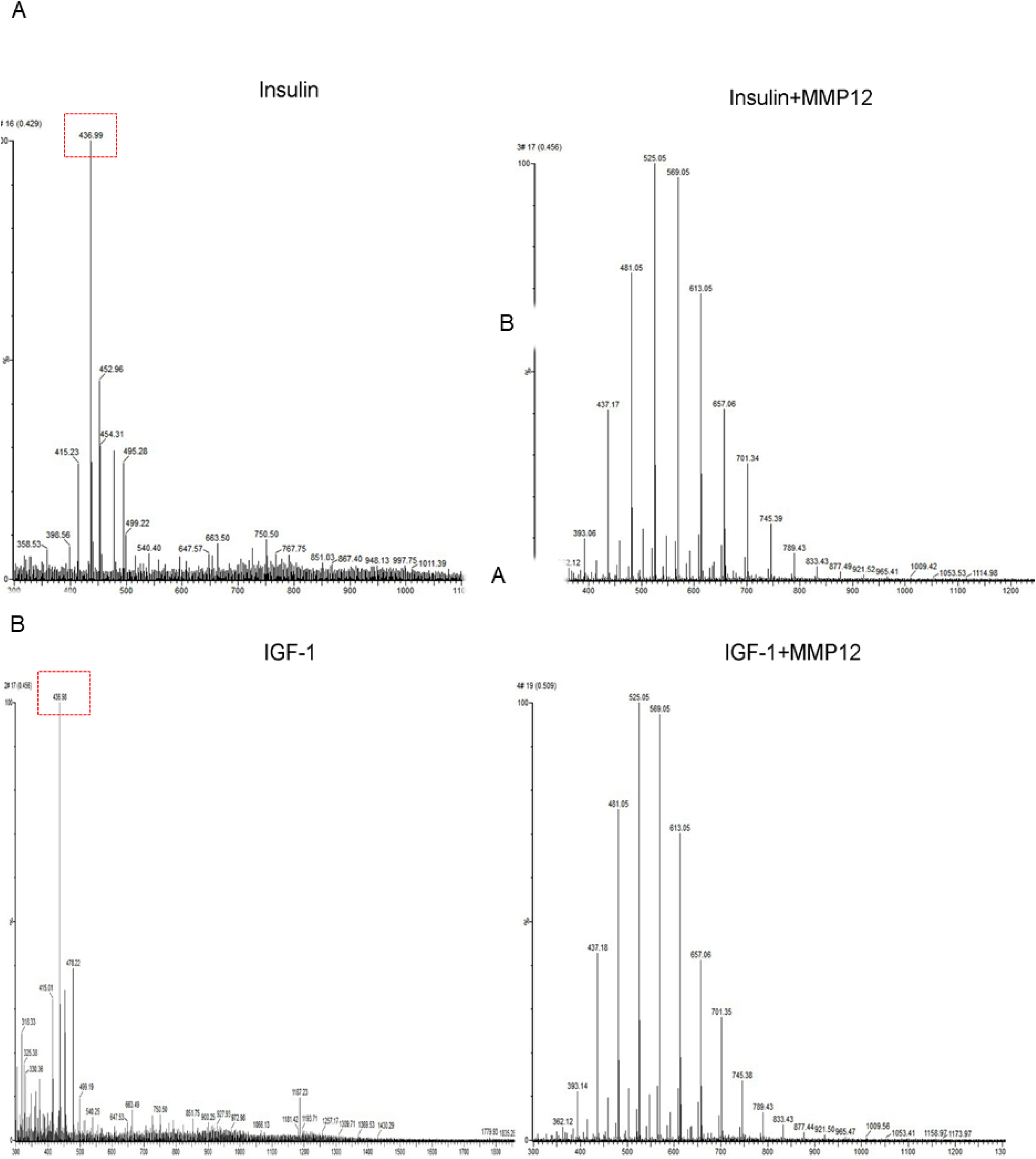
Ionization mass spectrometry analysis report: the characteristic peak of insulin (insulin-like growth factor 1) disappears after incubating with MMP12. (A) The characteristic peak of insulin (m/z = 436.99) and disappeared after incubation with MMP12. (B) The characteristic peak of insulin-like growth factor 1 (m/z = 436.98) disappeared after incubation with MMP12.

**Figure S3.**
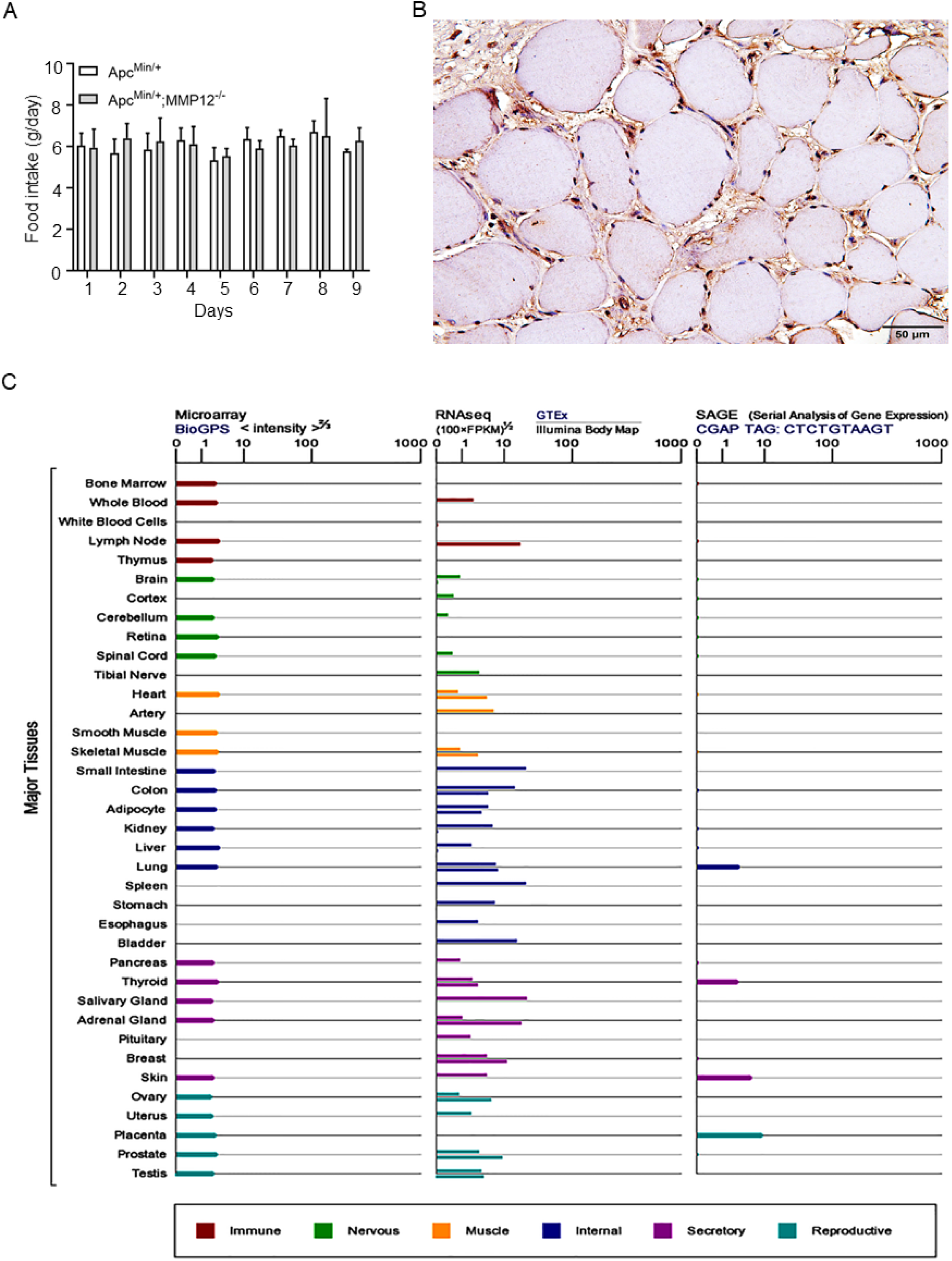
Knockout of MMP12 does not affect the food intake of Apc^Min/+^ mice and MMP12 is expressed in bone marrow, muscle, liver and adipose tissue. (A)There was no difference in food intake between Apc^Min/+^ mice and Apc^Min/+^; MMP12^-/-^ mice. The weight of food consumed after fasting for 8 h was measured every day starting on 17-week. Each group of mice had three cages, and each cage had 5 mice. (B) Immunostaining of MMP12-positive in muscle (gastrocnemius) from the clinical individual. Scale bar, 50μm. (C) A mRNA analysis of the data got from The Cancer Genome Atlas (TCGA) for GTEx, Illumina, BioGPS and SAGE of MMP12 gene in normal human tissues.

**Figure S4.**
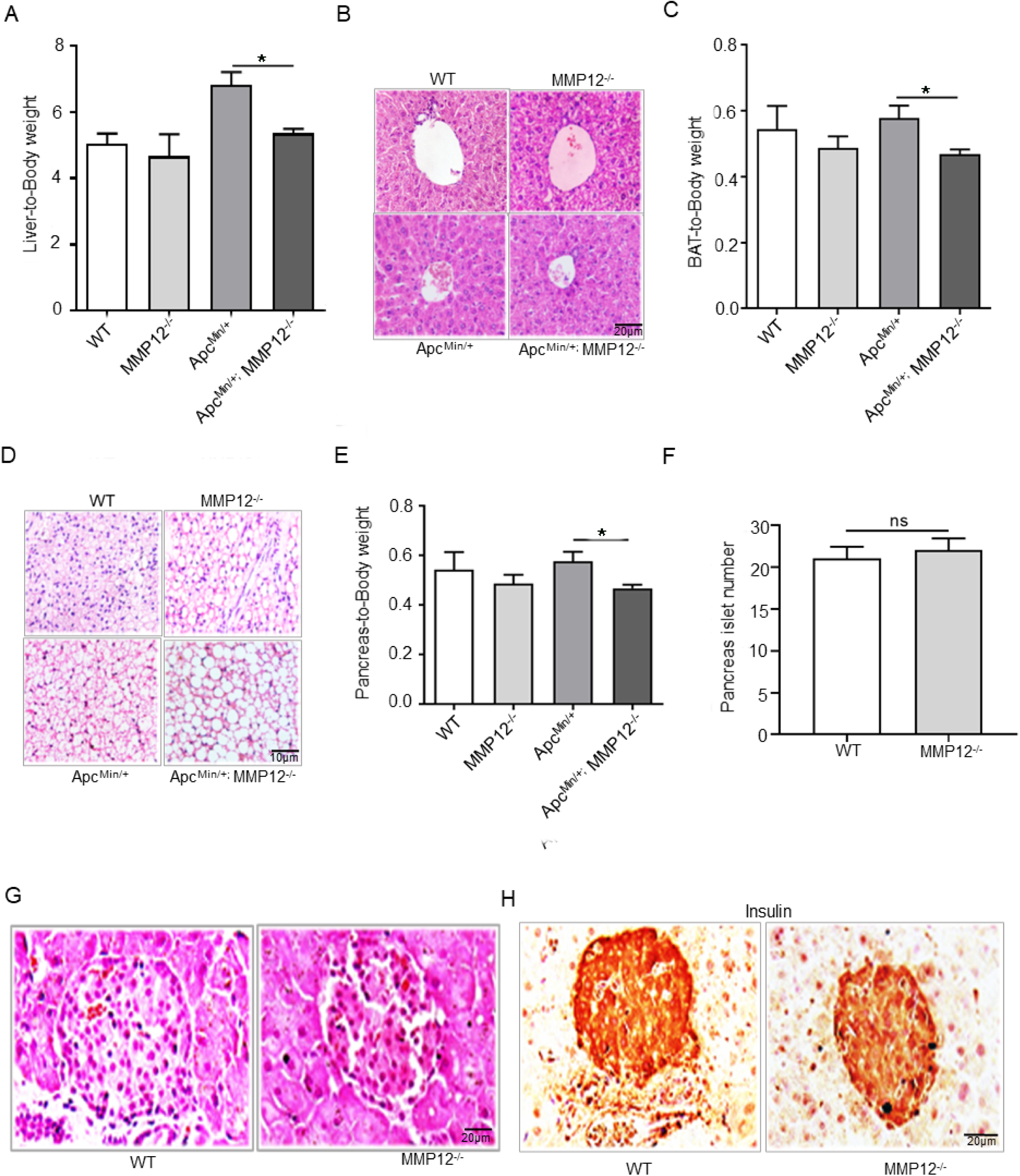
The effect of knocking out MMP12 in Apc^Min/+^ mice on liver, brown adipose tissue and pancreatic islets. (A)The liver-to-body weight ratio. (B) Hematoxylin and eosin staining (H&E) of the liver at 24 weeks. Scale bar, 20μm (\**P* < 0.05 data are shown as means ± SD; n = 6 per group). (C)Brown adipose tissue-to-body weight ratio was higher in Apc^Min/+^; MMP12^-/-^ mice than in Apc^Min/+^mice (\**P* < 0.05 data are shown as means ± SD; n = 6 per group). (D) The results of H&E indicate that white fat increased in brown fat in MMP12 knockout mice. Scale bar, 10μm. (E) The pancreas-to-body weight ratio (\**P* < 0.05 data are shown as means ± SD; n = 6 per group). (F) The number of islets between WT and MMP12^-/-^ mice (*P*> 0.05; data are shown as the means ± SD; n = 6 per group). (G) The staining results of H&E for pancreas WT and MMP12^-/-^ mice at 24 weeks. Scale bar,20μm. (H)Immunostaining of insulin-positive in WT mice and in MMP12^-/-^ pancreases at 24 weeks of age. Scale bar, 20μm.

**Figure S5.**
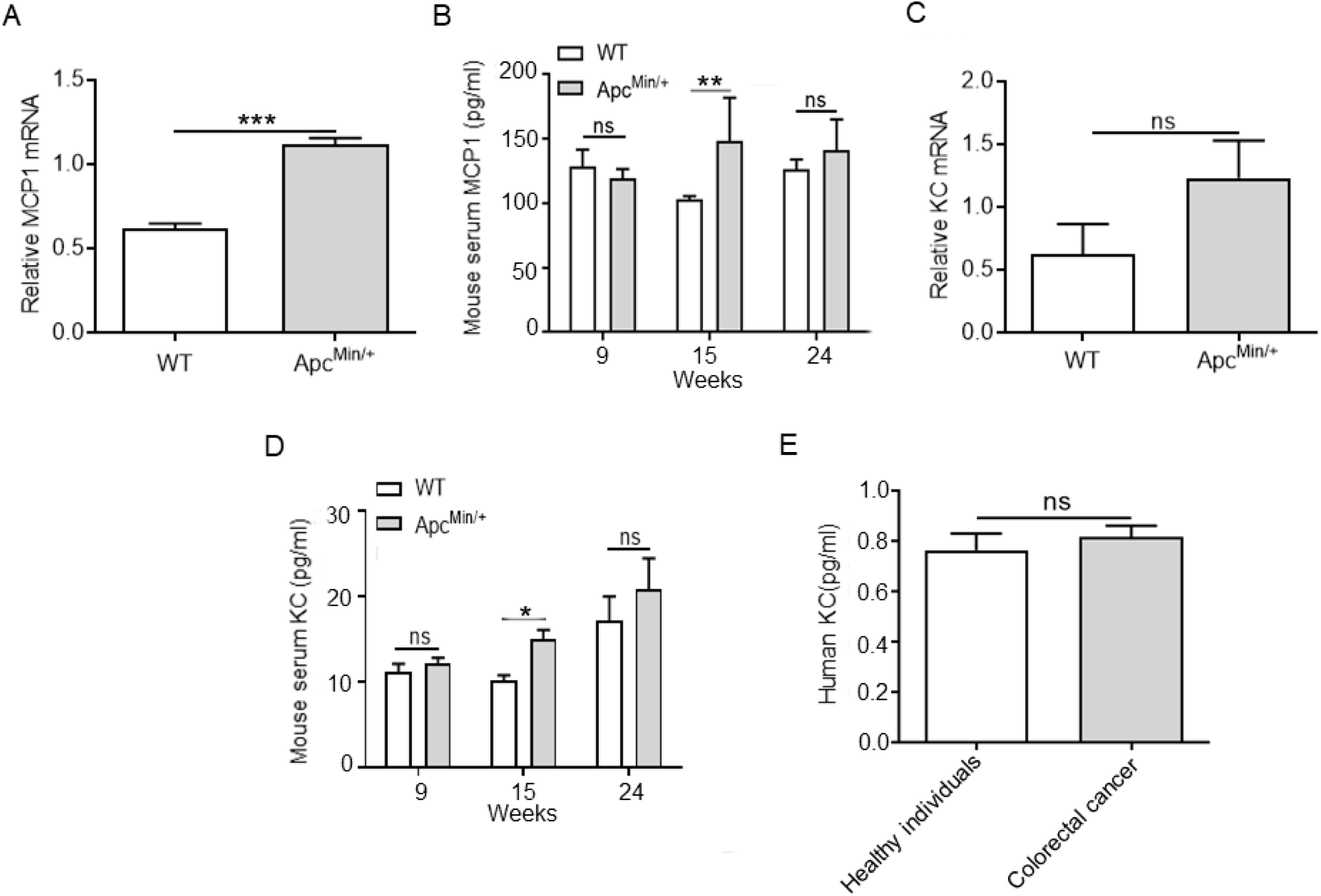
Serum monocyte chemoattractant protein 1(MCP1) and keratinocyte-de-rived chemokine(KC) did not change of Apc^Min/+^ mice in cancer cachexia at 24 weeks of age. (A, C) The mRNA expression of MCP1 and KC was validated in normal intestinal epithelium isolated from WT mice versus that in intestinal tumors isolated from Apc^Min/+^mice by quantitative PCR (\*\*\**P<*0.001 data are shown as means ± SD; n = 4 per group). (B, D) Serum MCP1 and KC in Apc^Min/+^ mice versus WT mice at 9, 15 and 24 weeks (\*\**P<* 0.01, \**P* < 0.05 data are shown as means ± SD; n = 6 per group). (E) Serum KC in normal healthy individuals and colorectal cancer patients by enzyme-linked immunosorbent assay (*P*> 0.05; data are shown as the means ± SD; n = 6 per group).

**Figure S6.**
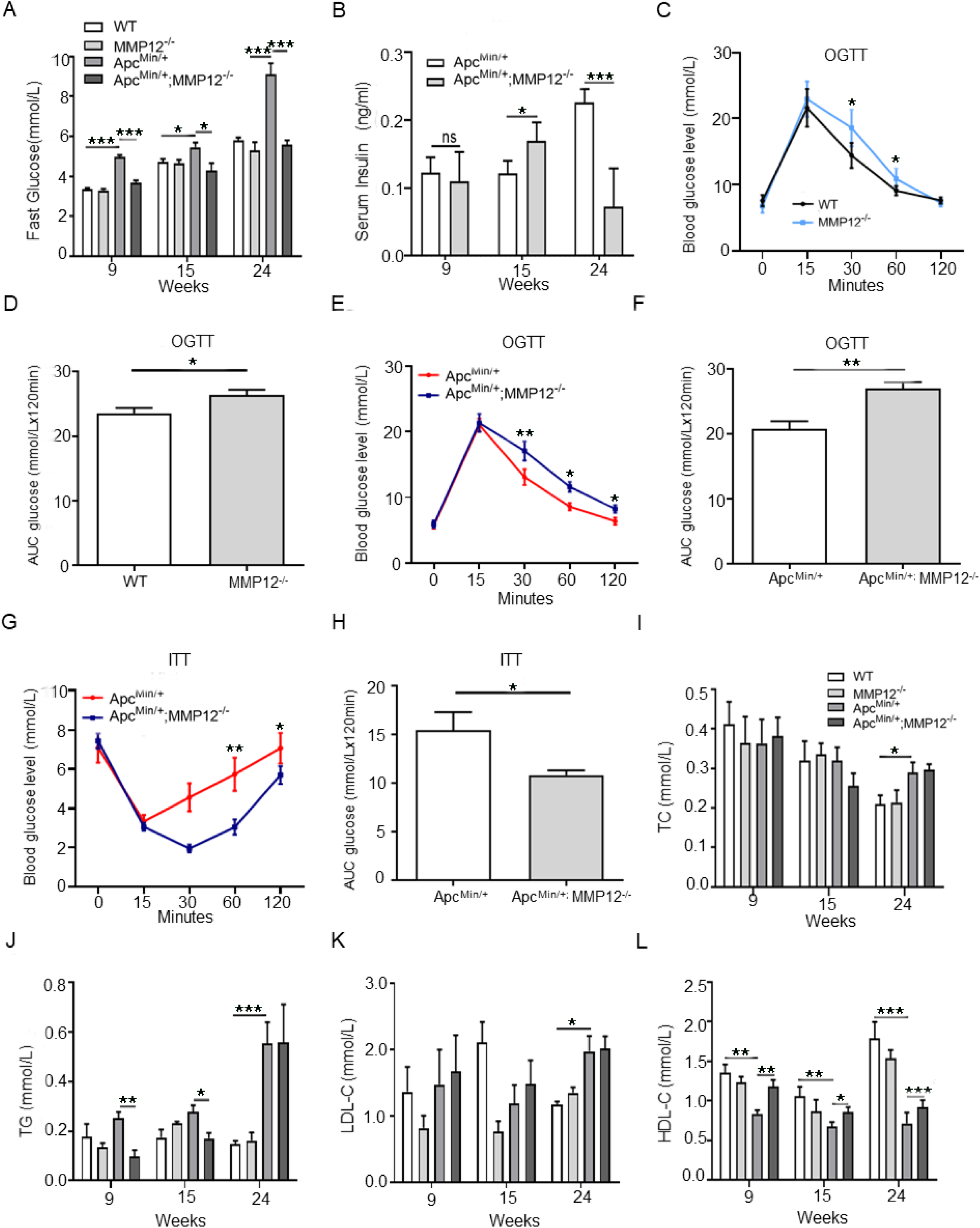
Knockout MMP12 affects glycolipid metabolism in Apc^Min/+^ mice. (A)Fasting plasma glucose levels at 9, 15 and 24 weeks of age in WT mice, MMP12^-/-^ mice, Apc^Min/+^ mice and Apc^Min/+^; MMP12^-/-^ mice (\*\*\**P<*0.001, \**P* < 0.05; data are shown as the means ± SD; n = 4 per group). (B) Enzyme-linked immunosorbent assay was used to detect fasting serum insulin levels in Apc^Min/+^and Apc^Min/+^; MMP12^-/-^ mice at approximately 9, 15 and 24 weeks of age (\*\*\**P<*0.001; \**P<* 0.05; data are shown as the means ± SD; n = 4 per group). (C) Oral Glucose Tolerance Test (OGTT): WT, MMP12^-/-^ mice were fasted for 4 h and then administered glucose (75 IU/kg), (\**P<* 0.05; data are shown as the means ± SD; n = 6 per group). (D) Area under the curves for the OGTT-AUG, which was increased in MMP12^-/-^ mice compared with that in WT mice (\**P* < 0.05 data are shown as means ± SD; n = 6 per group). (E) OGTT: Apc^Min/+^ and Apc^Min/+^; MMP12^-/-^ mice were fasted for 4 h and then administered glucose (75 IU/kg), (\*\**P<* 0.01, \**P* < 0.05 data are shown as means ± SD; n = 6 per group). (F) Areas under the curves for the OGTT-AUG, which was significantly increased in Apc^Min/+^; MMP12^-/-^ mice compared with that in Apc^Min/+^mice (\*\**P<* 0.01, data are shown as means ± SD; n = 6 per group) (G) Insulin tolerance test (ITT): mice were fasted for 4 hours; Apc^Min/+^ and Apc^Min/+^; MMP12^-/-^mice then received an ip injection of insulin (\*\**P<* 0.01, data are shown as means ± SD; n = 6 per group). (H) Areas under the curves for the ITT-AUG, which was significantly decreased in Apc^Min/+^; MMP12^-/-^ mice compared with that in Apc^Min/+^mice (\**P* < 0.05 data are shown as means ± SD; n = 6 per group). (I-L) Quantitative determination of serum total cholesterol (TC), total triglyceride (TG), low density lipoprotein-cholesterol (LDL-C), high density lipoprotein-cholesterol (HDL-C) in WT mice, MMP12^-/-^ mice, Apc^Min/+^ mice and Apc^Min/+^; MMP12^-/-^ mice by kits at 9, 15 and 24 weeks of age (\*\*\**P<*0.001,

**Figure S7.**
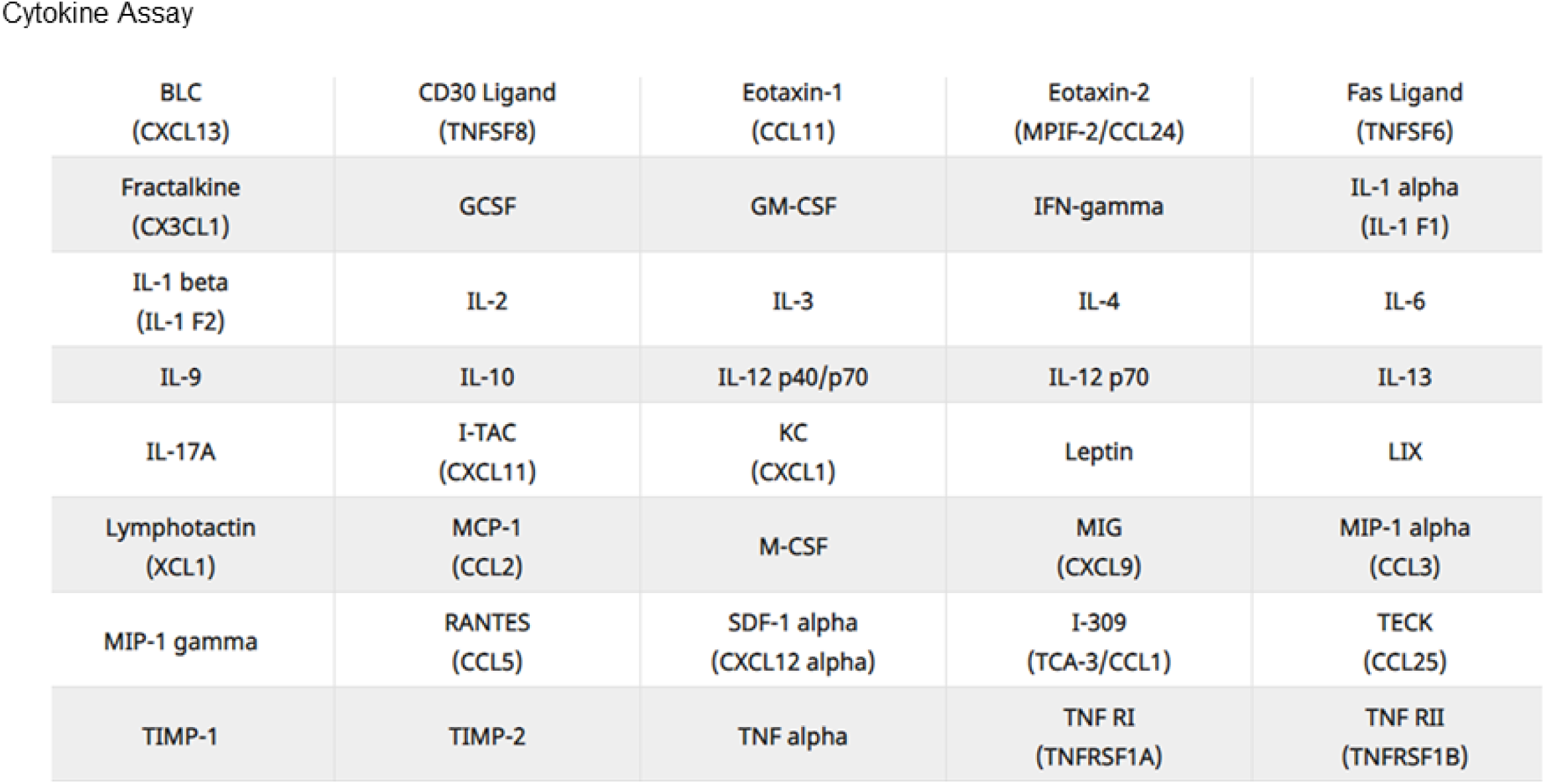
Specific detection factors in the protein chip. Protein chip contains 40 kinds of cytokine detection indicators.

